# A conserved fungal Knr4/Smi1 protein is vital for maintaining cell wall integrity and host plant pathogenesis

**DOI:** 10.1101/2024.05.31.596832

**Authors:** Erika Kroll, Carlos Bayon, Jason Rudd, Victoria Armer, Anjana Magaji- Umashankar, Ryan Ames, Martin Urban, Neil A. Brown, Kim Hammond-Kosack

## Abstract

Filamentous plant pathogenic fungi pose significant threats to global food security, particularly through diseases like Fusarium Head Blight (FHB) and Septoria Tritici Blotch (STB) which affects cereals. With mounting challenges in fungal control and increasing restrictions on fungicide use due to environmental concerns, there is an urgent need for innovative control strategies. Here, we present a comprehensive analysis of the stage-specific infection process of *Fusarium graminearum* in wheat spikes by generating a dual weighted gene co-expression network (WGCN). Notably, the network contained a mycotoxin-enriched fungal module that exhibited a significant correlation with a detoxification gene-enriched wheat module. This correlation in gene expression was validated through quantitative PCR.

By examining a fungal module with genes highly expressed during early symptomless infection, we identified a gene encoding FgKnr4, a protein containing a Knr4/Smi1 disordered domain. Through comprehensive analysis, we confirmed the pivotal role of FgKnr4 in various biological processes, including morphogenesis, growth, cell wall stress tolerance, and pathogenicity. Further studies confirmed the observed phenotypes are partially due to the involvement of FgKnr4 in regulating the fungal cell wall integrity pathway by modulating the phosphorylation of the MAP-kinase MGV1. Orthologues of *FgKnr4* are widespread across the fungal kingdom but are absent in other Eukaryotes, suggesting the protein has potential as a promising intervention target. Encouragingly, the restricted growth and highly reduced virulence phenotypes observed for *ΔFgknr4* were replicated upon deletion of the orthologous gene in the wheat fungal pathogen *Zymoseptoria tritici*. Overall, this study demonstrates the utility of an integrated network-level analytical approach to pinpoint genes of high interest to pathogenesis and disease control.

## Introduction

The wheat crop (*Triticum* species) plays a crucial role in global food security, contributing about 20% of dietary calories and protein worldwide (Saldivar, 2016), while also supplying essential nutrients and bioactive food components (Shewry and Hey, 2015). Pathogen and pest burden substantially contribute to wheat losses globally, accounting for ∼21.5% of wheat losses annually (Savary et al., 2019). Of these, the five highest global contributors to wheat yield and quality losses are all fungal diseases and include Fusarium Head Blight disease (FHB) and Septoria tritici blotch disease (STB), which account for 2.85% and 2.44% of wheat losses, respectively (Savary et al., 2019).

FHB is a mycotoxigenic pre-harvest fungal disease of most cereals, caused by different *Fusaria* within the *Fusarium sambucinum* species complex that is increasingly prevalent in most cereal growing regions globally (O’Donnell et al., 2000; Kanja et al., 2021; Johns et al., 2022; Armer et al., 2024). Floral Infections lead to contamination of grain with mycotoxins that are subject to strict legal limits in different global regions (European Commission, 2006; EFSA, 2017; AHDB, 2023). Despite ongoing endeavours to manage FHB, mycotoxin contamination continues to significantly impact the economies of cereal and livestock producers, as well as the food, drink, and feed industries (Latham et al., 2023). The B-type sesquiterpenoid deoxynivalenol (DON) is the most common FHB mycotoxin in European food and feed wheat (Johns et al., 2022). The globally predominant DON producing species is *Fusarium graminearum* (O’Donnell et al., 2000). During wheat spike colonisation, *F. graminearum* undergoes a biphasic mode of infection. Initially, the fungus evades the host immune response by growing between cells, causing no visible symptoms for ∼3 days. This is followed by an extended symptomatic stage marked by wheat tissue bleaching and reduced grain development behind the advancing hyphal front (Brown et al., 2010, 2011). STB disease on wheat leaves is caused by the fungus *Zymoseptoria tritici.* This fungus has an extended symptomless stage of infection ∼9 days, followed by a switch to symptomatic disease (Goodwin et al., 2011; Steinberg, 2015). However, unlike *F. graminearum*, *Z. tritici* colonisation is strictly confined to the sub-stomatal cavities and apoplastic spaces, without ever invading host cells (Kema et al., 1996). Both pathogens are currently managed using semi effective sources of host resistance mediated by major genes or QTLs (Brown et al., 2015; Bai et al., 2018; Buerstmayr et al., 2020) as well as fungicide applications (Fones and Gurr, 2015; Torriani et al., 2015; Buerstmayr et al., 2020; Kanja et al., 2021). But effective control faces escalating issues caused by fungicide resistance (Estep et al., 2015; Lucas et al., 2015; McDonald et al., 2019; de Chaves et al., 2022). There is a critical need to develop new methods to combat these and other wheat fungal pathogens.

Understanding the genetic and molecular mechanisms driving host infection in numerous interaction types continues to be a major goal of the international molecular plant pathology community (Nelson, 2020; Jeger et al., 2021). Gene expression data can be organised into co-expression networks, which group genes based on shared co-expression patterns. Network representations are advantageous because these present biological data on a systems-wide level, clustering genes in modules representative of specific stages or functions. This modelling can be achieved using the weighted gene co-expression network analysis (WGCNA) framework (Langfelder and Horvath, 2008). WGCNA has been repeatedly applied to analyse fungal gene expression data. For instance, this approach has been employed to identify effectors in *Magnaporthe oryzae* (Yan et al., 2023), shared genes during *Fusarium oxysporum* infection across multiple hosts (Cai et al., 2022), and virulence genes of *Colletotrichum siamense* (Liu et al., 2023). Although WGCNA has been used to study wheat host responses to *F. graminearum* infection (Kugler et al., 2013; Pan et al., 2018) and responses of *F. graminearum* under *in vitro* stress (L. Zhang et al., 2022; Park et al., 2023), there has been no study of wheat-*F. graminearum* co-expression profiles during infection.

To gain deeper insight on the expression patterns of genes during the different stages of the *F. graminearum* infection the WGCNA framework was used to generate a fungal pathogen/wheat dual co-expression network. Significantly, this framework can facilitate the correlation of both fungal and host plant expression (Mateus et al., 2019). Within this approach, genes are grouped into modules based on shared co-expression patterns separately for the pathogen and the host. Modules are then correlated between the pathogen and host networks to predict shared expression dynamics. In this study, correlated expression between a mycotoxin gene-enriched fungal module and a detoxification gene-enriched wheat module, validated the host-pathogen network. The study then focused on the unique fungal module F16, characterised by high expression levels during the earliest symptomless infection stage, and led to the discovery of *FgKnr4*. A subsequent comprehensive experimental analysis revealed the pivotal role of *FgKnr4* in various biological processes, including morphogenesis, growth, cell wall stress tolerance, and virulence in *F. graminearum*. The *Knr4* gene is not restricted to *F. graminearum* but is distributed widely across the fungal kingdom and is absent in other Eukaryotes. The various mutant phenotypes observed in the *F. graminearum ΔFgknr4* strain were replicated upon deletion of the orthologous gene in the wheat pathogen *Z. tritici*. Overall, this study highlights the value of using network analyses to model spatio-temporal pathogen-host interactions and to identify novel conserved genes associated with virulence.

## Results

### Generation of a dual *F. graminearum*-wheat co-expression network

*F. graminearum* floral infections can be divided into symptomatic or symptomless stages of infection. Disease spread through the rachis internodes (RI) can be further broken down to four different key stages of infection. Namely early symptomless (RI7-8), late symptomless (RI5-6), early symptomatic (RI3-4), and late symptomatic (RI1-2) infection **(Figure 1A)**. A spatio-temporal transcriptomics dataset of *F. graminearum* floral infection of the susceptible wheat cultivar Bobwhite, which distinguishes between these key distinct stages, was previously generated (Dilks et al., 2019). This dataset also included spikelet tissue (SP) sampled at 3 (early symptomatic) and 7 days post-infection (dpi) (late symptomatic). The WGCNA framework (Zhang and Horvath, 2005; Langfelder and Horvath, 2008) was used to construct a dual co-expression network to model fungal pathogen/crop interaction in wheat using this dataset.

**Figure 1.**
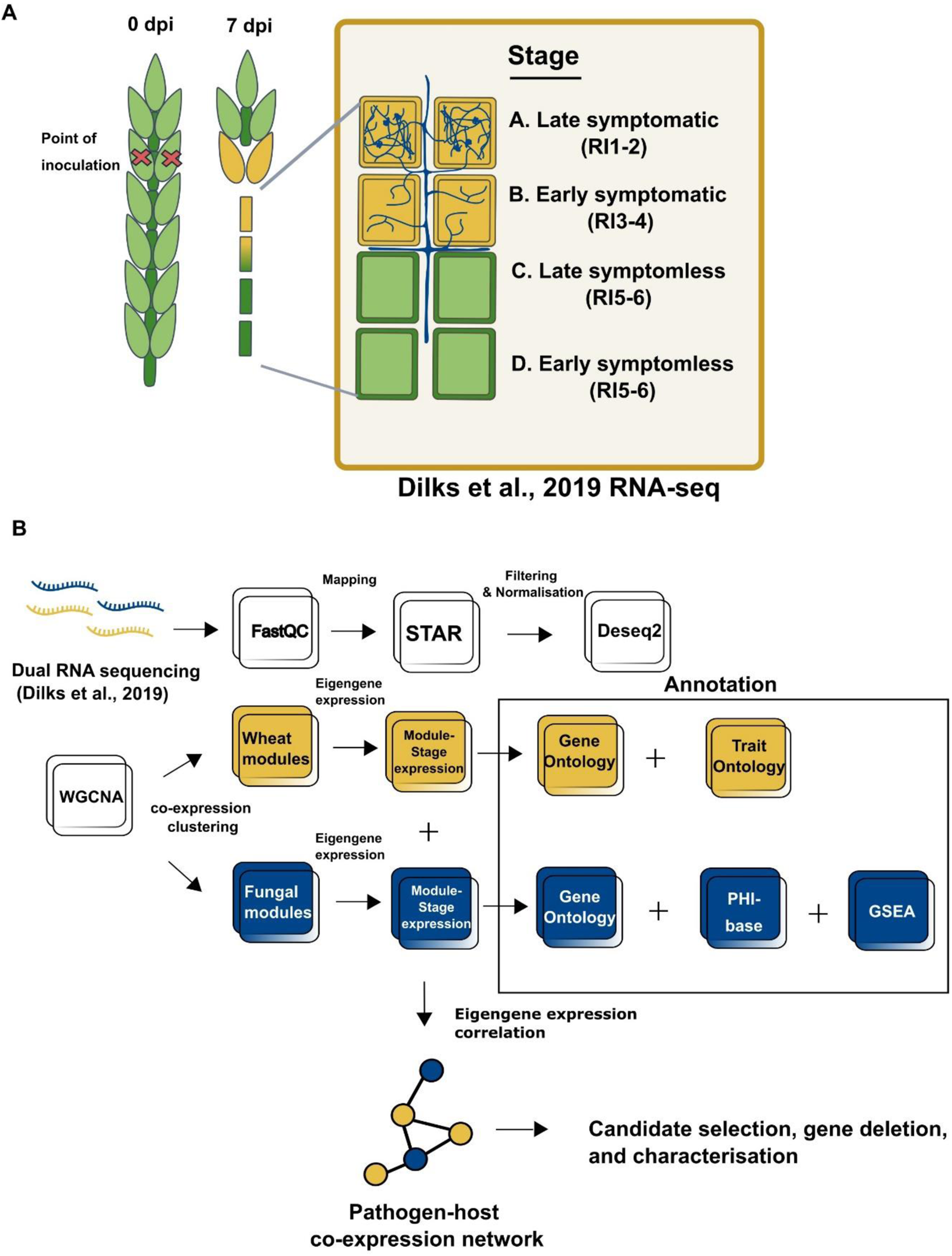
Dual RNA-seq dataset and bioinformatics pipeline used for constructing the dual co-expression network. **A.** Schematic illustration depicting the symptomatic (yellow) and symptomless (green) stages of *Fusarium graminearum* infection of wheat spikes denoted as stages A through D, corresponding to tissue samples collected for generating the RNA-seq data published in Dilks et al. 2019. *F. graminearum* hyphae growing in either the apoplast or inside the wheat cells are depicted in blue. **B.** Summary outlining the bioinformatics pipeline used for processing raw reads and constructing the dual RNA-seq network. The dual RNA-seq reads were initially processed together (processes depicted as white squares) before being separated to generate two distinct weighted gene co-expression networks. The bioinformatic pipelines are annotated accordingly, with yellow indicating the wheat reads-only pipeline and blue indicating the fungal reads-only pipeline. Annotation includes Gene Ontology terms (GO), Trait Ontology terms (TO), unique Gene Set Enrichment Analysis (GSEA), and PHI-base phenotypes. The modules from the two separate networks are then correlated to each other by their Eigengene values to form the dual co-expression network.

Normalised counts were used to generate two distinct networks: one for *F. graminearum* and another for *T. aestivum*. The *F. graminearum* network consisted of 10,189 genes organised into 18 modules (with 2629 – 60 genes per module), while the *T. aestivum* network consisted of 47,458 genes distributed among 25 modules (with 23063 – 83 genes per module) **(Figure 2 – figure supplement 1, Supplementary File 1)**. Both networks met scale free model criteria at their selected soft thresholding power **(Figure 2 – figure supplement 2 A-B)**. The examination of module quality statistics found that each module within both networks were of a high quality (Z-Summary > 10), with the exception of F16 (Z-Summary = 9.67), which still markedly surpasses the minimum Z-Summary score of > 2 (Langfelder et al., 2011) **(Figure 2 – figure supplement 2C**). This indicates a substantial preservation of modules compared to a random selection of all network genes. Additionally, preservation statistic calculations confirmed that all modules maintain preservation (Z-summary > 2) across both networks with all modules of the wheat network and the majority of the fungal modules (11/18) having strong preservation (Z-summary > 10) **(Figure 2 – figure supplement 2D**). These findings suggest a consistent preservation of within-network topology across modules (Langfelder et al., 2011). For each module, a single summarised expression pattern, the eigengene value, was calculated. The fungal and wheat modules were correlated by their eigengene expression values, and modules displaying significant correlation (*p* ≥ 0.001) formed the dual co-expression network **(Figure 2A)**.

**Figure 2.**
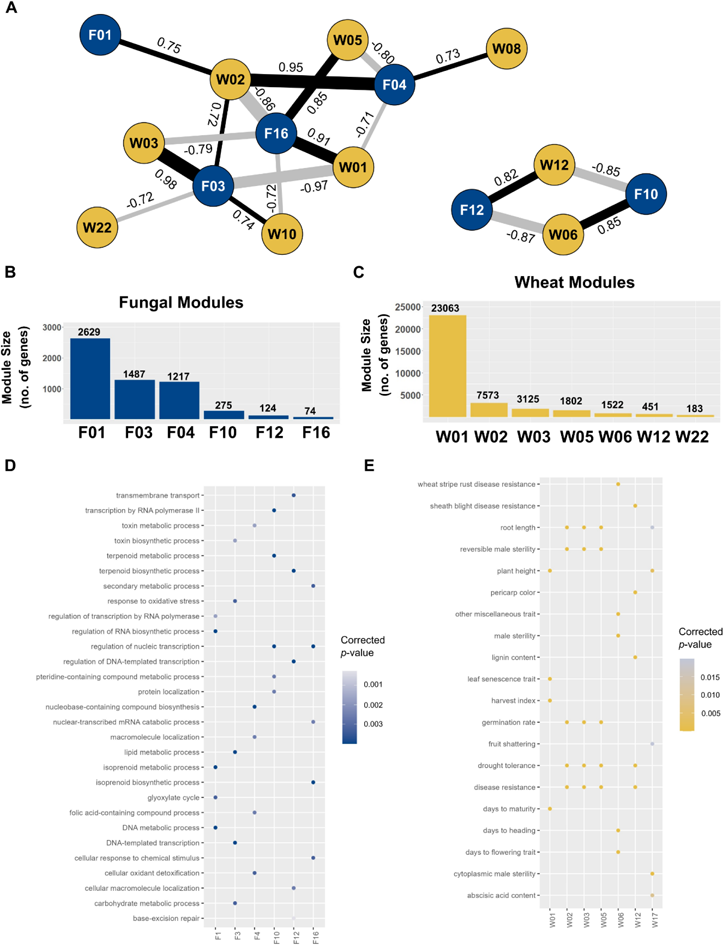
Dual Fungal-Wheat co-expression network. **A.** Network summarising significant co-expression patterns (*p* ≥ 0.001) between fungal modules (blue nodes) and wheat modules (yellow nodes). Positive correlations are depicted as black edges, while negative correlations are shown as grey edges. R-squared values are indicated next to edges, with edge width corresponding to the value. **B.** Fungal modules sizes. **C.** Wheat module sizes (**Supplementary File 1**). **D. Fungal module enrichment.** Significant (*p* ≤ 0.05) Biological Processes (BP) Gene Ontology (GO) enrichment results for all fungal modules in the network. Higher significance is indicated by darker blues. **E. Wheat module enrichment.** Significant Plant Trait Ontology (TO) enrichment results (*p* ≤ 0.05) for all wheat modules in the network. Higher significance is indicated by brighter yellows.

To gain insight into the function of individual modules, a Gene Ontology (GO) enrichment analysis was performed for both network sets (**Figure 2D-E**, **Figure 2 – figure supplement 1)**. To confirm these enrichment patterns were not due to chance, a random network was generated for both the fungal and wheat datasets. No significant enrichment was found for the random wheat network and fungal network.

Among the eight wheat modules within the dual co-expression network, five of them were significantly enriched for disease resistance genes (TO:0000112, *p ≥*0.05) and one was specifically enriched for wheat stripe rust resistance genes (TO:0020055) **(Figure 2E)**, suggesting the wheat modules in the network are needed for plant defence. One of these wheat modules, W12, was significantly enriched in the GO terms detoxification (GO:0098754; *p* = 7.13 x 10^-7^) and response to toxic substances (GO:0009636; *p* = 2.11 x 10^-6^). This module was correlated to the fungal module F12, which was enriched in genes belonging to the trichothecene biosynthesis (*TRI)* gene cluster (*p* = 1.92 × 10^-4^) and for the GO term terpenoid biosynthesis (GO:0016114 ; *p* = 0.00085) **(Figure 2D**, **Table 1)**. Notably, the module F12 was most highly expressed in the late symptomless stage of infection.

**Table 1.** Function of correlated expression between wheat and fungal modules. This table illustrates the relationship between wheat and fungal gene expression at different stages of infection, detailing the associated functions and key fungal genes involved.

| Expression Stages | Wheat Module | Predicted function | Correlated Fungal Module | Key Fungal Genes | Fungal Gene Functions | References |
| --- | --- | --- | --- | --- | --- | --- |
| Early symptomless stage of infection | W01 | Maintenance genes (photosynthesis, RNA modification) and early defence response. | F16 | *FgNPC1* | Regulation of membrane trafficking and sterol metabolism, which are essential for maintaining cellular integrity and function during the infection stages. | Breakspear et al. 2011 |
|  |  |  |  | *Gzc2h045* | Msn2 C2H2 transcription factor, associated with virulence and coordination of adaptation to environmental stressors including heat, osmotic, and oxidative stress. | Son et al., 2011; John et al., 2021 |
|  | W05 | Disease resistance genes, including reactive oxygen species genes associated with programmed cell death response to restrict pathogen spread. |  | *GzCon7* | Msn2 C2H2 transcription factor, associated with virulence and regulation of cell wall biosynthesis. | Son et al., 2011; John et al., 2021 |
|  |  |  |  | *FgSrp2* | Pre-mRNA processing, alternative splicing, and virulence | Zhang et al., 2020 |
| **Early symptomless and late symptomatic stages of infection** | W06 | Enriched in protein catabolism and autophagy, involved in immune signalling, programmed cell death, and necrotrophic damage control. | F10 | *KP4L-1, KP4L-2, KP4L-3* | Necessary for virulence, provide competitive advantage during new niche occupation, essential for intraspecific interactions at high fungal density | Lu and Faris, 2019; Vicente et al., 2022 |
|  |  |  |  | *FgOS-2, FgAtf-1* | Regulation of secondary metabolite production, sexual reproduction, and stress tolerance | Nguyen et al., 2013 |
|  |  |  |  | *FgHyd3, FgHyd5* | Attachment to hydrophobic surfaces, production of aerial mycelia | Shin et al., 2022 |
| **Late symptomless to early symptomatic** | W12 | Detoxification, response to toxic substances, and defence response. | F12 | *TRI* genes (*TRI3, TRI4, TRI11, TRI12, TRI14*) | Production of DON mycotoxin needed for virulence. | Dyer et al., 2005; Kimura et al., 2007 |

Expression of this module then rapidly decreases during the symptomatic stages of infection. Module F12 therefore appears to be positioned specifically at the transition between the late symptomless stage and the early symptomatic stage. The production of the DON mycotoxin is essential for the transition to the extensive symptomatic stage (Cuzick et al., 2008; Jansen et al., 2005). DON inhibits protein translation, which then eventually leads to cell death and the bleached phenotype distinctive of symptomatic *F. graminearum* infection (Desmond et al., 2008; Arunachalam and Doohan, 2013). High expression of module F12 in the symptomless stage is also supported by previous data which found that genes involved in mycotoxin biosynthesis are highly expressed in symptomless wheat tissue (Brown et al., 2017). The correlation with the wheat module W12 therefore implies that detoxification genes in the module are being expressed in response to production of fungal mycotoxins.

Interestingly, the fungal module F10 contains genes that are highly expressed in the earliest and latest stages of *F. graminearum* infection, but not intermediate stages **(Figure 4).** The fungal module F10 includes the Killer toxin 4 genes (*KP4L*) -1, -2, and -3. These genes also have some of the highest module membership scores (>0.90) within the module. The *KP4L* genes are necessary for virulence and expressed during both self and non-self interactions **(Table 1)**. It is suggested that KPL4 proteins provide *F. graminearum* with a competitive advantage when occupying new niches (Vicente et al., 2022), which would explain their expression during the earliest stage of infection. High expression during late infection may be necessary for intraspecific interactions, when the fungus is coordinating growth at a high fungal density.

The stress-responsive mitogen-activated protein kinase *FgOS-2* is a key regulator *in F. graminearum* and acts upstream of the ATF/CREB-activating transcription factor *FgAtf-1* **(Table 1)**. Both *FgOS-2* and *FgAtf-1* cluster in module F10. These proteins are involved in broad functions, including secondary metabolite production, sexual reproduction, and stress tolerance (Nguyen et al., 2013). Module F10 also contains two hydrophobin genes, *FgHyd3* and *FgHyd5*. *FgHyd3* is necessary for attachment to hydrophobic surfaces, while both genes are necessary for the production of aerial mycelia **(Table 1)**. These genes are likely to play a crucial role during early infection for surface attachment and are possibly expressed again during the late stage of infection to facilitate the production of aerial mycelia.

The fungal module F10 is correlated with the wheat module W06 (R = 0.85, *p* = 6 x 10^-6^), which is enriched in protein catabolism (GO:0010498; *p* = 1.60 x 10^-19^) and autophagy (GO:0006914; *p =* 2.31 x 10^-4^) **(Table 1)**. Autophagy plays a dual role in plant immunity where it is involved in immune signalling and programmed cell death to restrict pathogen spread, but also in response to pathogen induced necrotic cell death (Sertsuvalkul et al., 2022). Therefore, it is likely these genes are expressed during early infection as an immediate immune response and then expressed again in highly colonised tissue for late-stage necrotrophic damage control.

### Wheat genes in module W12 are expressed in response to DON production

To validate the correlation between modules F12 and W12 **(Figure 3A)**, expression of wheat genes in the detoxification module W12 in response to *F. graminearum* infection without DON was examined. This was achieved by inoculating wheat plants with either the wild-type *F. graminearum* reference strain PH-1, or the DON deficient *ΔFgtri5* mutant strain generated in the PH-1 background. Expression of three wheat genes was studied, including two phenylalanine ammonia-lyases (*PAL1* and *2*; TraesCS4A02G401300 and TraesCS2D02G377200) which were annotated with the term disease resistance (TO:0000112), and a predicted transmembrane exporter, detoxification gene 16 (*DTX16*; TraesCS5B02G371100).

**Figure 3.**
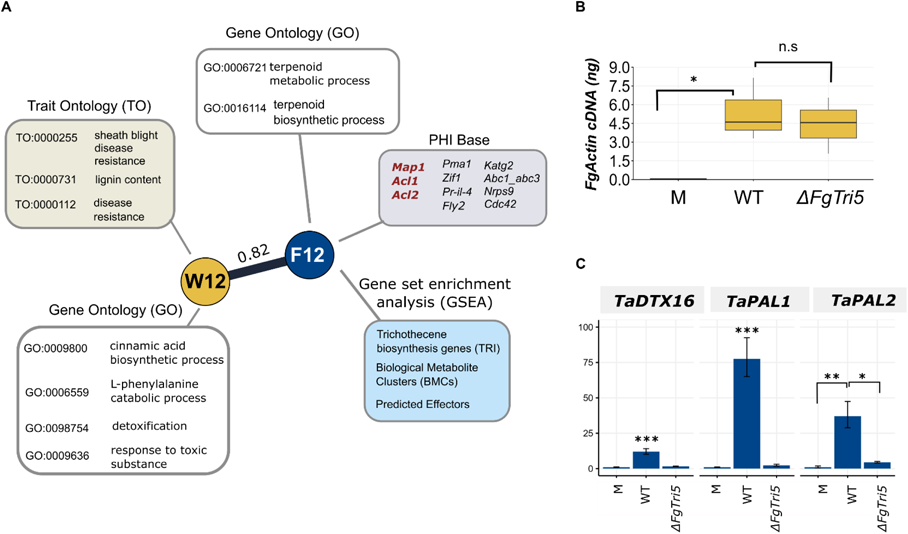
Validation of correlation between the trichothecene mycotoxin biosynthesis gene enriched module (F12) and the detoxification gene enriched module (W12). A. Modules F12 (N = 124) and W12 (N = 451) depicted with significant enrichment annotations and genes with known phenotypes from PHI-base. Three genes listed in red in the PHI-base annotation (grey box) exhibit a loss of pathogenicity phenotype, while the remaining genes display a reduced virulence phenotype when individually deleted in *F. graminearum*. B. Equal levels of fungal burden were observed in tissue samples (*p* > 0.05). Absolute quantity of actin cDNA in Mock, *ΔFgtri5*, and wild-type (WT)-recovered RI1-2 tissue sampled at 3 dpi. Significance was determined by a one-way ANOVA followed by Tukey HSD correction. C. Normalised fold change expression of selected W12 wheat genes in Mock, *ΔFgtri5*, and WT-recovered RI1-2 tissue sampled at 3 dpi (N = 3). Significance is denoted as * = *p* ≤ 0.05, ** = *p* ≤ 0.01, and *** = *p* ≤ 0.001. Significance was determined by a one-way ANOVA followed by Tukey HSD correction.

The first two rachis internodes below the point of inoculation (POI) were sampled at 3 days post inoculation (dpi). Levels of *FgActin* cDNA were not significantly different between treatments **(Figure 3B)**. Expression of the three wheat genes from module W12 was significantly lower in the *ΔFgtri5* infected samples relative to wild-type infection **(Figure 3C)**. This indicates that expression of genes in module W12 is correlated with DON production, thereby supporting the correlated co-expression patterns observed between modules of the two networks.

### Dual co-expression networks as a tool to identify key genes necessary for virulence

To pinpoint *F. graminearum* genes that are necessary for virulence, the stage specific expression patterns of each module was examined **(Figure 4**, **Figure 4 – figure supplement 1**). The module F16 is uniquely highly expressed during the earliest stages of infection, with markedly decreased expression at all the other stages of infection. This module is highly correlated to two wheat modules. These are W01 (R = 0.91; *p* = 5 x 10^-7^) and W05 (R = 0.85, *p* = 2 x 10^-5^). W01 is the largest wheat module and is enriched for defence response genes (GO:0006952; *p* = 3.60 x 10^-08^), but also maintenance genes which include photosynthesis (GO:0015979; *p* = 4.59 x 10^-29^) and RNA modification (GO:0009451; *p* = 1.42 x 10^-47^) GO terms. The wheat module W05 is enriched for disease resistance (TO:0000112, *p* = 2.55 x 10^-178^), suggesting that despite the continued symptomless infection the host is already expressing genes for defence. Four genes in module F16 result in reduced virulence when individually deleted. These are *FgNPC1* (sterol trafficking) (Breakspear et al., 2011), *FgSrp2* (mRNA splicing) (Zhang et al., 2020), and the transcription factors *Gzcon7* and *Gzc2h045* (Son et al., 2011) **(Table 1, Supplementary File 2)**. However, no gene deletion mutants exhibiting a loss of pathogenicity or highly reduced virulence phenotype have yet been identified within this module, even though the eigengene expression pattern clearly indicates an association with the early establishment of the fungus in this key host tissue.

**Figure 4.**
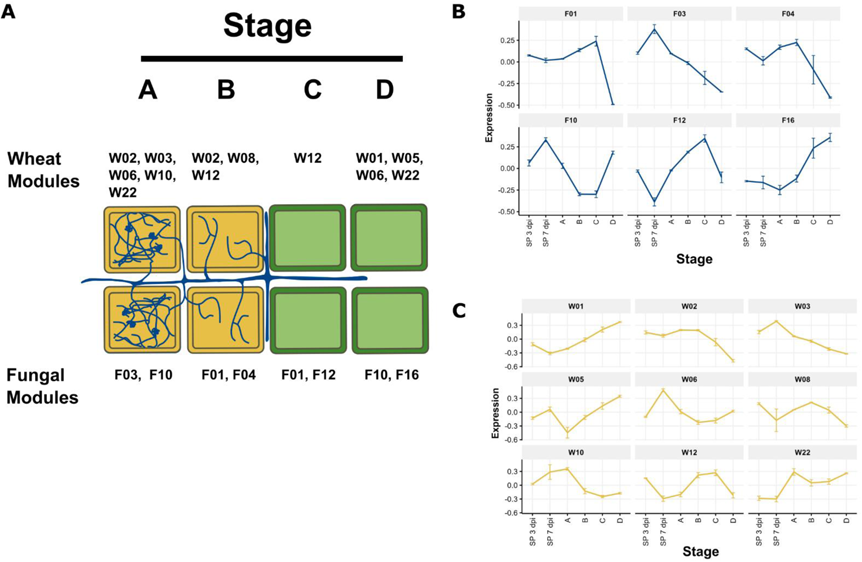
Stage-specific expression of modules in the dual co-expression network. **A.** Expression of modules across stages of *F. graminearum* infection. Illustration depicting symptomatic (yellow) and symptomless (green) stages of infection (A through D) annotated with specific modules (W or F) from the dual co-expression network that were highly expressed at specific stages **B. Eigengene summarised expression of fungal modules** and **C. wheat modules**. Eigengene summarised expression plots illustrating the expression patterns of genes in wheat modules across different stages of infection as illustrated in panel A, along with spikelet tissue (SP) at 3 and 7 dpi.

To identify genes in F16 that are likely involved in virulence, the 74 genes within this module were examined. Key genes were defined as those exhibiting elevated module membership (MM) within the module, which were also strongly correlated (R > |0.70|) with corresponding wheat modules. Genes with a high MM value have expression patterns closely aligned with the module’s overall eigengene expression and are the most representative of the module. The initial candidate gene list was selected by starting with the 15 key genes with the highest MM within the module. Genes were then excluded that were likely to have functional redundancy (i.e. belonged to a gene family or had ancient paralogues within PH-1) to avoid compensatory effects when performing single gene deletion **(Supplementary Table S1)**. Ultimately, only two genes met these criteria: FGRAMPH1_0T23707 and FGRAMPH1_01T27545. FGRAMPH1_01T27545 has been previously characterised as the Niemann–Pick type C gene (*FgNPC1*). *FgNPC1* is necessary for sterol trafficking, with its deletion resulting in ergosterol accumulation within the vacuole and a reduced virulence upon wheat infection (Breakspear et al., 2011). Orthologue analysis identified that the FGRAMPH1_0T23707 gene was a 1:1 orthologue of Killer-nine resistant 4 (KNR4) in *Saccharomyces cerevisiae* (Martin et al., 1999), therefore the orthologue in *F. graminearum* is henceforth referred to as *FgKnr4*.

### *FgKnr4,* a key gene of module F16, is necessary for establishment of fungal infection

*FgKnr4* was deleted using a split hygromycin replacement cassette **(Figure 6 – figure supplement 1 A-B)**. *T. aestivum* cv. Bobwhite was inoculated at anthesis with three independent *ΔFgknr4* transformants. No symptomatic disease progression past the inoculated spikelets was observed with each *ΔFgknr4* transformant **(Figure 5 A-B)**. While the inoculated spikelets developed symptoms, these did not exhibit full bleaching of the spikelet characteristic of FHB infection. Instead, eye-shaped lesions formed akin to those evident following *ΔFgtri5* mutant infection **(Figure 5C)** (Cuzick et al., 2008). Plating of surface sterilised wheat dissected into its constituent parts revealed the absence of fungal growth in un-inoculated spikelets **(Figure 5 – figure supplement 1**). Nevertheless, browning was noted in the rachis tissue immediately adjacent to the point inoculated spikelet, accompanied by fungal growth. However, this colonisation did not occur past the rachis internode of the 3rd spikelet. These data suggest that, despite entering the rachis, the *ΔFgknr4* mutant is unable to grow through the rachis node tissue and re-enter other spikelets. Microscopic examination revealed a more pronounced plant defence response to *ΔFgknr4* infection. This was characterised by a visibly reduced fungal burden **(Figure 5D)**.

**Figure 5.**
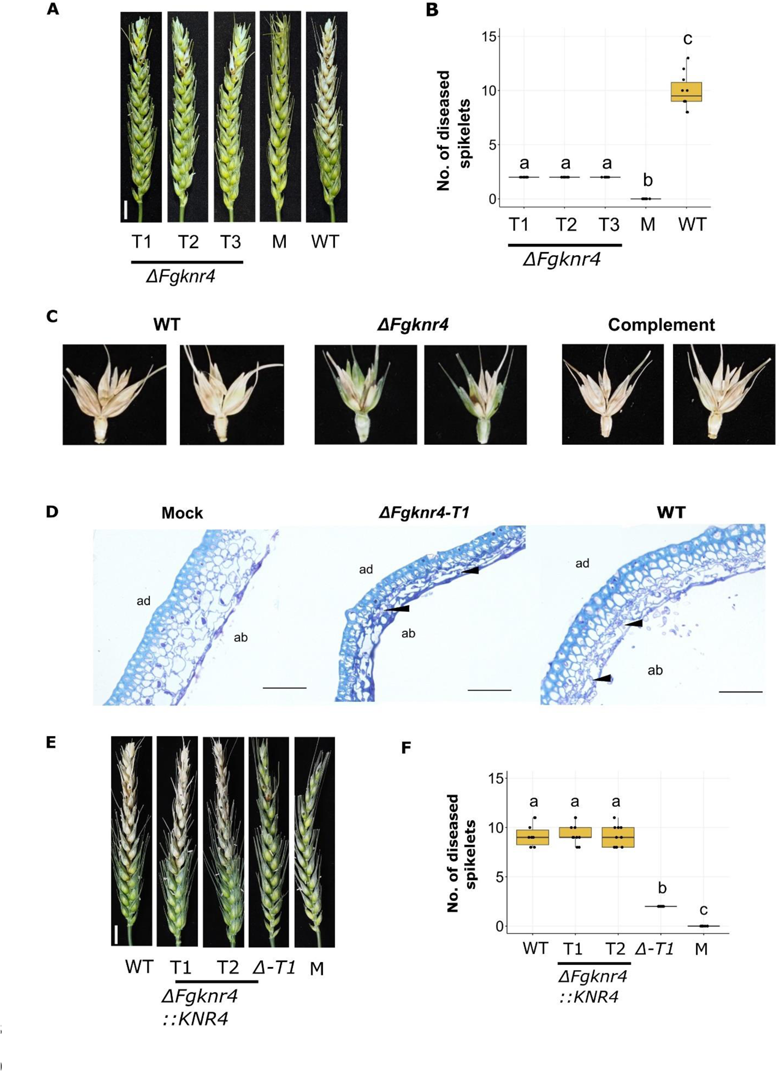
Decreased virulence observed during *in planta* infection with *ΔFgknr4*. **A.** Wheat spike infection assay done on the susceptible cultivar Bobwhite point inoculated with sterile water only (Mock), wild-type *F. graminearum* conidia, or conidia from three independent single gene deletion *F. graminearum* mutants lacking *Knr4* (*ΔFgknr4,* T1-3). Images were captured at 15 dpi. Scale bar = 1 cm. **B.** Number of diseased spikelets per wheat spike at 15 dpi. Letters indicate significant differences (ANOVA, TukeyHSD *p* < 0.05). C. Symptom development on the inoculated spikelets and adjacent rachis tissues at 15 dpi D. Ultra-thin 1µm LR White resin sections stained with 0.1% Toluidine Blue for visualisation of wheat cell walls (light blue) and fungal hyphae (purple). Black arrows indicate fungal hyphae. Fungal hyphae typically proliferate in the abaxial cell layer. Ab = abaxial and ad = adaxial. Scale bar = 50 µm. Tissue harvested at 7 dpi. **E.** Wheat spike infection complementation assay done on the susceptible cultivar Bobwhite treated with conidia either from wild-type *F. graminearum*, different complemented transformants (*ΔFgknr4::KNR4-T1* and *T2*), the single gene deletion mutant (*ΔFgknr4-T1*), or sterile water (mock). Images were taken at 15 days post inoculation. **F.** Number of diseased spikelets per wheat spike at 15 dpi. Letters indicate significant differences (ANOVA, TukeyHSD *p* < 0.05).

**Figure 6.**
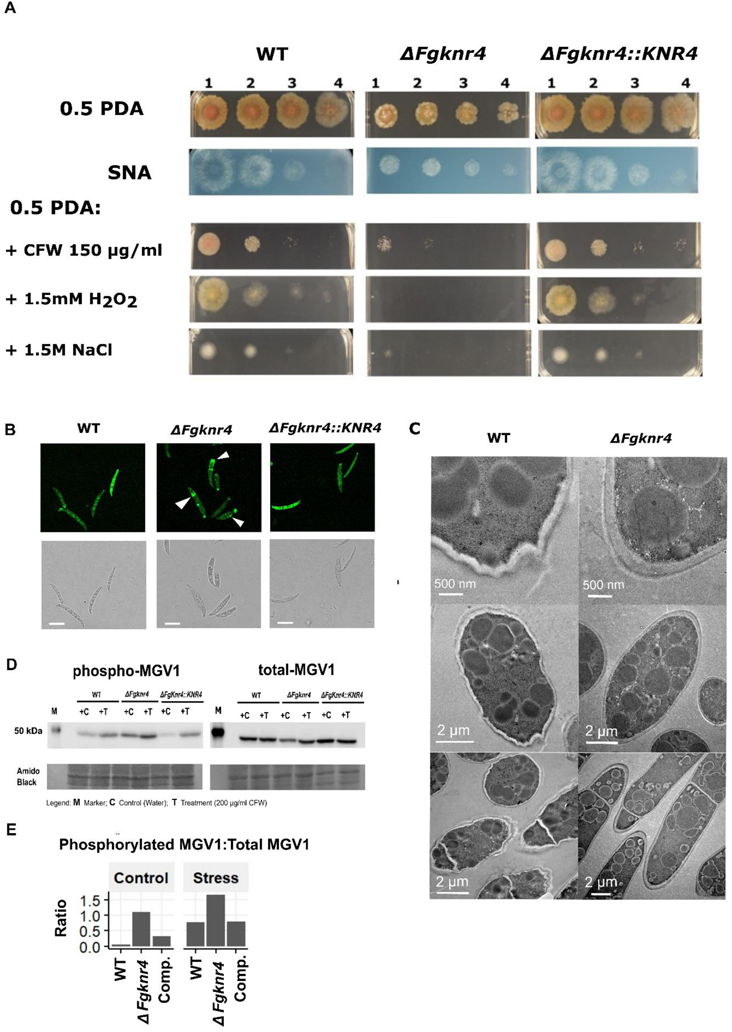
Cell wall stress sensitivity and abnormal cell wall morphology of *ΔFgknr4*. **A**. Dilution series of wild-type (WT), *ΔFgknr4*, and *ΔFgknr4::KNR4* strains on Synthetic Nutrient Agar (SNA) and half-strength Potato Dextrose Agar (0.5 PDA) with and without the addition of calcofluor white (CFW), hydrogen peroxide (H_2_O_2_), and sodium chloride (NaCL). The dilution series begins at 1: 1 x 10^6^ and continues with 10-fold dilutions (2: 1/10, 3: 1/100, and 4: 1/100). Images taken after 3 days of growth at room temperature. **B**. Abnormal chitin deposition patterns in *ΔFgknr4* conidia. Chitin-stained in conidia visualised using Wheat Germ Agglutinin Alexa Fluor™ 488 Conjugate (WGA). Scale bar = 50 µm. **C**. TEM imaging of wild-type and *ΔFgknr4* conidia, showing differences in cell wall structure **D.** Western blot of proteins extracted from, *ΔFgknr4* and *ΔFgknr4::KNR4* mycelium incubated with (T) or without (C) the addition of 200 µg/ml Calcofluor White (CFW) for 24 h. Phospho-p44/42 MAPK (Erk1/2) and p44/42 MAPK (Erk1/2) antibodies were used to detect phosphorylated and total MGV1, respectively. Amido black total protein staining was performed to compare protein loading. E. Ratio of phosphorylated MAPK/total MAPK based on quantification of band intensity.

Despite highly reduced virulence, DON mycotoxin was detected in the inoculated spikelet and attached rachis internodes (≥ 0.2 ppm). However, DON was undetectable in the neighbouring uninoculated spikelet (< 0.2 ppm). Complementation of the mutant with wild-type *FgKnr4* restored virulence to wild-type levels **(Figure 5 E-F)**.

### *FgKnr4* influences cell wall structure, stress resistance, and growth

*In vitro* growth of *ΔFgknr4* was examined by culturing the fungus on both high or low nutrient agar. In both conditions a decreased growth rate relative to the wild-type was apparent **(Figure 6A**, **Figure 6 – figure supplement 1 and 2**). In addition to this, conidia of *ΔFgknr4* appear smaller than wild-type **(Figure 6 – figure supplement 3 A,C)**. Despite these morphological differences *ΔFgknr4* retains the ability to produce perithecia and ascospores, albeit 8 days later than the wild-type **(Figure 6 – figure supplement 3 D-E)**.

Stresses encountered by the fungus during *in planta* infection were mimicked *in vitro* using chemical stressors. *ΔFgknr4* had increased susceptibility to osmotic stress (1.5M NaCl), oxidative stress (H_2_O_2_), and calcofluor white induced cell wall damage compared to the wild-type and complemented strains **(Figure 6A**, **Figure 6 – figure supplement 1C & 2C)**.

These susceptibilities may be due to changes in the cell wall structure of the *ΔFgknr4* strain. Corroborating this hypothesis, staining for chitin found an irregular deposition of chitin on the *ΔFgknr4* conidial cell wall, specifically along the tips and septa of the conidia **(Figure 6B**, **Figure 6 – figure supplement 4**). Furthermore, an irregular cell wall structure was observed upon transmission electron microscopy (TEM) analysis of the *ΔFgknr4* conidia, indicative of an abnormal cell wall composition **(Figure 6C**, **Figure 6 – figure supplement 5**).

The *FgKnr4* (F16) module was correlated with the wheat module W05, which exhibits a significant enrichment in the term oxidative stress (TO: 0002657; *p* = 3.88 x10^-34^) that encompasses a total of 1143 genes. Among these genes are two respiratory burst oxidase homologues (RBOH), specifically a predicted homolog of RBOF (TraesCS1A02G347700) and RBOHE (TraesCS5D02G222100), along with predicted catalase homologues, CAT3 (TraesCS7B02G473400) (Ghorbel et al., 2023; Yan Zhang et al., 2022), and two CAT4 genes (TraesCS5B02G023300, TraesCS5D03G0079400) (Andleeb et al., 2022). W05 is also enriched for sodium content (TO: 0000608; *p* = 0.00014) and salt tolerance (TO: 0006001; *p =* 3.00 x 10^-18^). The necessity of a functional *FgKnr4* gene in oxidative and osmotic stress tolerance **(Figure 6A**, **Figure 6 – figure supplement 1C & 2C**) suggests that *FgKnr4* is critical during this early infection stage, where the fungus confronts hydrogen peroxide and osmotic stress induced by the plant.

The involvement of *FgKnr4* in cell wall metabolism was further studied by examining its effect on the cell wall integrity pathway (CWI). The fungal CWI pathway is triggered in response to various stresses (e.g. oxidative stress, osmotic pressure, cell wall damage) (Dichtl et al., 2016) and in *F. graminearum* is activated through the phosphorylation of the MAP-kinase (MAPK) *FgMGV1* (Hou et al., 2002; Yun et al., 2014). A Western blot was run on mycelium samples grown with and without a cell wall stress (calcofluor white). Constitutive activation of MGV1 in the absence of stress and increased phosphorylation under stress was observed in *ΔFgknr4* when compared to the wild-type (**Figure 6D-E**). This finding is consistent with previous observations in *S. cerevisiae* (Martin-Yken et al., 2003). This reinforces the biological function of *FgKnr4*, suggesting an involvement in fungal stress responses and cell wall morphology in *F. graminearum*.

### The orthologous gene in wheat pathogen *Zymoseptoria tritici* is also important for cell wall integrity and virulence on wheat

Analysis of the Knr4 protein conservation found that orthologues were highly distributed across the Dikarya, occurring in both Ascomycota and Basidiomycota **(Figure 7).** Notably, no orthologues of the gene were found in other Eukaryotes, highlighting its specificity to the fungal lifestyle. This high level of conservation across fungi suggests that phenotypes observed in *F. graminearum* may also be conserved in other economically significant pathogenic fungi.

**Figure 7.**
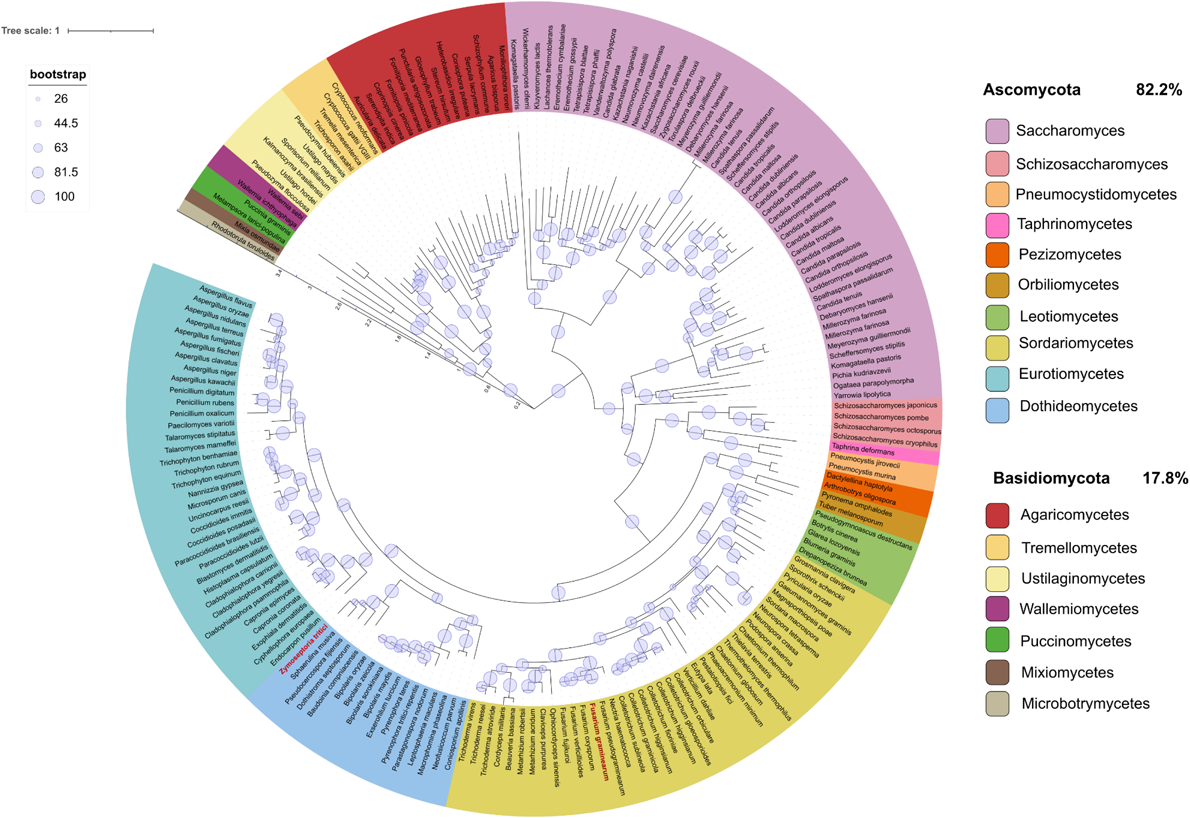
Distribution of KNR4 orthologues across eukaryotes reveals exclusive presence in fungi. A phylogenetic tree depicting the distribution of *KNR4* orthologues across Eukaryota, with the positions of *F. graminearum* and *Z. tritici* highlighted in red. Different taxonomic levels are indicated in various colours as specified in the legend, alongside the percent distribution of orthologues between Ascomycota and Basidiomycota. Evolutionary distances between species or taxa are denoted by an internal scale (range 0 - 3.5) Bootstrapping confidence values are depicted as pale blue circles, with increasing size corresponding to higher confidence levels.

The orthologous *KNR4* gene in another wheat fungal pathogen *Z. tritici* (*ZtKnr4*, Mycgr3G105330) was disrupted to test for conserved gene function. Despite the phylogenetic distance between the two fungi, the *FgKnr4* and *ZtKnr4* proteins share 43.5% pairwise identity. Mirroring the phenotype observed in *F. graminearum,* reduced virulence (chlorosis but limited to no necrosis) was observed when wheat leaves were inoculated with *ΔZtknr4* **(Figure 8A)**. In addition to this the *ΔZtknr4* mutant was susceptible to calcofluor white induced cell wall stress and exhibited reduced hyphal branching **(Figure 8B-C)**. These results highlight the potential of employing the *Fusarium*-wheat dual co-expression approach to gain insights into fungal-plant interactions, both within *Fusarium* species and across the fungal kingdom.

**Figure 8.**
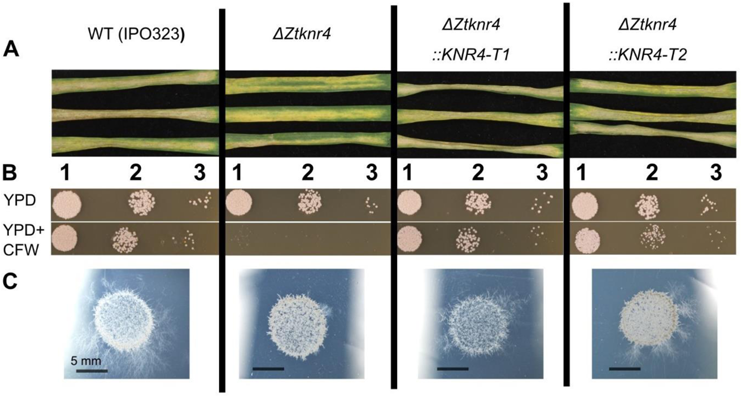
Functional characterisation of the *Zymoseptoria tritici ΔZtknr4* gene deletion mutant. **A.** Detached wheat leaves inoculated with wild-type (WT) *Z. tritici* (IPO323), *ΔZtknr4* mutant strain, and two complemented strains (*ΔZtknr4::KNR4-T1 and T2*). Image taken at 20 dpi. **B** WT *Z. tritici*, the *ΔZtknr4* mutant and two complemented strains (*ΔZtknr4::KNR4-T1 and T2*) spot inoculated onto YPD agar with (bottom) and without (top) calcofluor white (CFW). Dilution series begins at 1: 1 x 10^5^ and continues with 10-fold dilutions (2: 1/10 and 3: 1/100). Images taken after 3 days of growth at room temperature (RT). **C** WT *Z. tritici*, the *ΔZtknr4* mutant and two complemented strains (*ΔZtknr4::KNR4-T1 and T2*) spot inoculated onto 1% Tap Water Agar (TWA). Images taken after 10 days of growth at room temperature (RT).

## Discussion

The generated dual *F. graminearum-*wheat co-expression network was successfully used to identify a gene necessary for virulence. By analysing stage-specific modules of infection, module F16 was identified, which exhibited high gene expression levels during the symptomless stage of FHB infection. Within this module, the gene *FgKnr4* was found to have a high module membership score, indicating its central role in the module. Experimental validation showed that *FgKnr4* is essential for responding to chemical compounds that induce cell wall stress, early establishment of *in planta* infection, and subsequent disease progression in wheat spikes. Similarly, the deletion of *Knr4* in another pathogenic species, namely *Z. tritici* resulted in a reduced virulence phenotype in leaves and displayed a comparable cell wall stress phenotype. This highlights the utility of pathogen-host co-expression network analysis in identifying conserved virulence genes across wheat fungal pathogens.

The predictions from the WGCNA were validated for the F12-W12 correlation through the experimental confirmation of the co-regulation of the Fusarium trichothecene mycotoxin and wheat detoxification genes during infection. For Fusarium, module F12 was of exceptionally high interest because of its positioning specifically at the transition between the late symptomless stage and the early symptomatic stage. For wheat genes in the correlated module W12, the studied genes included two phenylalanine ammonia-lyases (*PAL1* and *2)* along with a predicted detoxifying efflux transporter (*TaDTX16*). Although *TaPAL1* and *2* have not been previously studied for their direct involvement in disease resistance in the wheat - Fusarium interaction, the *PAL* gene family is known to be associated with disease resistance and other phenotypes (Duba et al., 2019). In multiple plant species (including *Arabidopsis,* pepper (*Capsicum annuum)*, and rice (*Oryzae sativa)*), *PAL* is induced in response to biotic and abiotic stresses, which includes pathogen induced stress, (Hahlbrock and Scheel, 1989; Kim and Hwang, 2014; Tonnessen et al., 2015; Chen et al., 2017) and in numerous genetically incompatible host-pathogen interactions mediated by cognate R-Avr proteins including responses to fungi (Maher et al., 1994; Ramaroson et al., 2022). *TaDTX16* is part of the multidrug and toxic compound extrusion (MATE) gene family and was named after its orthologue in *Arabidopsis thaliana* (Li et al., 2002). DTX/MATE take part in heavy metal and lethal compound detoxification in plants and could be involved in mycotoxin detoxification (Perincherry et al., 2019). Previously, a wheat DTX gene was reported to be highly expressed in resistant cultivars of wheat compared to a susceptible wheat cultivar when infected with *F. graminearum* (Pan et al., 2018)*. TaDTX16* is located on chromosome 5BL within an interval harbouring a resistance QTL for defence against the necrotrophic fungal disease Septoria nodorum blotch (Li et al., 2021).

The characterisation of *FgKnr4*, underscores the importance of identifying genes necessary for full virulence through gene expression studies. This approach is essential because predicting the pathogenic potential of Fusarium species based solely on comparative genomics is challenging due to the absence of significant differences in secreted effector proteins, carbohydrate-active enzymes, or gene repertoires between pathogenic and endophytic strains of Fusarium and Fusarioid species (Hill et al., 2022). *FgKnr4* was investigated further for its multifaceted roles in growth, stress response, and cell wall integrity. Supporting previous findings in *Fusarium asiaticum* (Yu Zhang et al., 2022), this study demonstrates that *FgKnr4* is involved in regulating growth rate, conidial spore morphology, and sensitivity to osmotic and oxidative stresses, as well as virulence and cell wall stress tolerance in *F. graminearum*. Moreover, this study establishes that KNR4 influences the well-studied cytoplasmically located Mgv1 cell wall integrity (CWI) MAPK pathway (Xu et al., 2022), resulting in visible abnormalities of the conidial cell wall **(Figure 6C**, **Figure 6 – figure supplement 5**). The cell wall integrity pathway in *F. graminearum* is well-characterised, with each MAP-kinase in the cascade having been identified, studied, and shown to have roles in virulence and/or asexual and sexual spore formation (Hou et al., 2002; Jenczmionka et al., 2003; Urban et al., 2003; Zheng et al., 2012). Through experimental validation, our findings reveal an additional layer of control within the *F. graminearum* cell wall integrity pathway mediated by *FgKnr4*. This discovery contributes to and further improves our understanding of the regulatory mechanisms governing cell wall integrity in *F. graminearum*. This new finding also, offers the first insights into the regulatory effects of KNR4 in a filamentous fungus. This additional knowledge aid the development of novel strategies to mitigate losses caused by FHB disease and DON contamination.

*Z. tritici* possesses one of the most expansive publicly available eukaryotic pangenomes, with approximately 42% of its genes categorised as accessory (Plissonneau et al., 2018). *ZtKnr4* is part of the core *Z. tritici* genome of the European pangenome (Chen et al., 2023) and designated within the core orthogroup OG0008320 within the global (Europe, Asia, North and South America, Australia and Africa) pangenome (Badet et al., 2020). Given the highly variable nature of accessory chromosomes in *Z. tritici*, the assignment of *ZtKnr4* to the core genome in two separate pangenomic analyses underscores its importance in fungal physiology. *ZtKnr4* is also expressed throughout the wheat infection process (Rudd et al., 2015). Disruption of the gene resulting in a reduced virulence phenotype reinforces the potential of *ZtKnr4* as a candidate target for fungicide development, emphasising its significance in combating *Z. tritici* infections and mitigating agricultural losses. Despite the ever present global importance of STB disease for many decades (Dean et al., 2012; Savary et al., 2019) *Z. tritici* has far fewer functionally characterised genes, with only 99 genes with a characterised phenotype within the Pathogen Host-Interactions database and only 50 of these genes associated with a loss in pathogenicity or reduced virulence (Urban et al., 2022; Cuzick et al., 2023). The reduced virulence phenotype observed in the *ZtKnr4* mutant therefore marks a valuable contribution to the characterisation of one of the >9000 core genes across the known *Z. tritici* pangenomes (Badet et al., 2020; Chen et al., 2023).

The high conservation and exclusivity of KNR4 within the fungal kingdom, combined with its absence in other eukaryotes and its conserved function across related species, suggest that KNR4 could be an ideal target for intervention. This could be achieved through the development of chemical fungicides that disrupt the protein’s function (Aamir et al., 2018) or through the application of RNA interference techniques (Cools and Hammond-Kosack, 2013; Machado et al., 2018; Mann et al., 2023). Stricter EU regulation of chemicals suitable for fungicide use in agricultural, medical and/or veterinary settings (European Commission, 2022), combined with significant losses in fungicide efficacy due to the evolution of pathogen populations means there is a pressing need to identify new target sites for control. Therefore, this research not only advances our understanding of fungal virulence mechanisms but also offers promising directions for the development of effective strategies for disease control in agriculture.

## Materials and Methods

### Gene co-expression network analysis

RNAseq reads from Dilks et al. (2019) were provided by Dr Neil Brown (European Nucleotide Archive: PRJEB75530). Read quality was assessed with FastQC v. 0.11.9 (Andrews, 2010). Reads were mapped to a combined Fusarium – wheat genome, consisting of v. 5 of the *Fusarium graminearum* PH-1 genome (King et al., 2017) and the high confidence (HC) transcripts of the v. 2.1 of the International Wheat Genome Sequencing Consortium (IWGSC) *Triticum aestivum* genome (Zhu et al., 2021). Genome indexing and read alignment were performed using STAR aligner 2.7.8a. Soft clipping was turned off to prevent reads incorrectly mapping to similar regions of the highly duplicated hexaploid wheat genome. Reads were filtered using the filterByExpr function part of the R package Edge R v.3.32.1 (Robinson et al., 2010). Counts were normalised separately for fungal and wheat reads by performing a variance stabilising transformation (VST) using the DESeq2 v 1.30.1 R package (Love et al., 2014) in R (v4.0. 2, https://www.r-project.org/).

The VST normalised counts were filtered to remove any excessive missing values using the function goodSamplesGenesMS in the WGCNA R package (Langfelder and Horvath, 2008). Standard methods were implemented to generate the network using the WGCNA R package, with the following parameters. A signed-hybrid network was constructed using the filtered counts. The soft thresholding power (β) was uniquely selected per network according to scale free model criteria (Zhang and Horvath, 2005), where β = 9 for the fungal network and β = 18 for the wheat network **(Figure 2 – figure supplement 2**). A deepSplit of 3 was paired with a standard cutheight of 0.25. A minimum module size of 50 was selected to minimise potential transcriptional noise when assigning modules using smaller datasets (Oldham, 2014; Walsh et al., 2016). The function multiSetMEs from the WGCNA package was used to calculate module eigengene expression. Module eigengenes with similar expression profiles were then merged.

Module quality and preservation was calculated using the function modulePreservation present in the WGCNA R package (Langfelder and Horvath, 2008; Langfelder et al., 2011). When calculating module preservation, the original wheat or fungal network was considered the reference network. Then 50 different test networks were created, each built upon randomly resampling (with replacement) a proportion of samples from the original dataset. The average preservation metrics (i.e. Z-score) between the original network and the 50 test networks was calculated for both the fungal and wheat networks.

### Module Enrichment an Annotation

Gene ontology (GO) annotations of the v. 5 PH-1 genome (GCA_900044135.1) were generated using Blast2GO v .5 (Götz et al., 2008). Enrichment was calculated using a background set of all genes present in the fungal network. GO annotations for the IWGSC v.2.1 genome were provided by Dr Keywan Hassani-Pak of the KnetMiner team (Hassani-Pak et al., 2021). This was generated by performing a BLASTx search on the NCBI nb database using DIAMOND v 2.0.13-GCC-11.2.0 (Buchfink et al., 2015), then Blast2GO v.5 was used to annotate the BLAST hits with GO terms. GO term enrichment was calculated for each high level GO ontology (Biological Process, Molecular Function and Cellular Component) using the R package topGO v 2.46.0 (Alexa and Rahnenfuhrer, 2009).

Plant Trait Ontology (TO) (Cooper et al., 2024) enrichment analysis was performed using annotations derived from the KnetMiner knowledge graph (release 51) for wheat (Hassani-Pak et al., 2021) and KnetMiner datasets and enrichment analysis notebooks are available at https://github.com/Rothamsted/knetgraphs-gene-traits/. Predicted effectors were determined using EffectorP v.3.0 (Sperschneider and Dodds, 2022). Alongside this, predictions to identify extracellularly localised genes were done using SignalP v6.0 (Teufel et al., 2022). Custom *F. graminearum* gene set enrichment of the network modules was calculated by performing a Fisher’s exact test using all the genes in the fungal network as the background gene set. A BH correction was calculated for both GO and custom enrichments (Benjamini and Hochberg, 1995). Modules were deemed significantly enriched if P-corr < 0.05.

Gene lists included in GSEA consisted of predicted secreted effector proteins, alongside known gene families associated with virulence, such as biological metabolite clusters (BMCs) (Sieber et al., 2014), polyketide synthases (Gaffoor et al., 2005), protein kinases (Wang et al., 2011) and transcription factors (Son et al., 2011). Due to their well-established importance in *F. graminearum* pathology, a separate enrichment for genes of the *TRI* gene cluster was also performed.

Annotation from PHI-base was obtained by mapping genes to version PHI-base (v4.16) annotation using UniProt gene IDs and any through Decypher Tera-Blast™ P (TimeLogic, Inc. Carlsbad, California, USA) (E-value = 0) against the PHI-base (v4.16) BLAST database (Cuzick et al., 2023).

### Fungal material and growth conditions

*F. graminearum* strains were cultured and conidia prepared as previously described (Brown et al., 2010). Fungal strains were grown for 4 days on nutrient-rich potato dextrose agar (PDA), nutrient-poor synthetic nutrient agar (SNA; 0.1% KH2PO4, 0.1% KNO3, 0.1% MgSO4·7H2O, 0.05% KCL, 0.02% glucose, 0.02% sucrose and 2% agar) and PDA with different cell wall stresses. Plates were point inoculated with 20 μl of 4-fold dilution series starting with 1 x 10^6^ conidia/ml. For the growth rate assay, fungi were grown on PDA and images were taken at 7 days. Surface sterilisation of wheat spikes was performed by submerging single wheat spikelets in 1/8 diluted thin bleach for 3 min, followed by three washes with distilled H2O. Dissection was done using a razor blade to separate the point inoculated spikelets and adjacent spikelets **(Figure 5 – supplement 1**). Wheat tissue was placed on SNA and images taken after 3-day incubation at room temperature in the dark. Perithecia induction was achieved as described in Cavinder et al. (2019). All plate images were taken using an Olympus OM-D camera using a 60mm ED M.Zuiko macro lens. Conidia and ascospore images were taken using the Axiomager 2 (Zeiss, Oberkochen, Germany) under brightfield illumination. Conidia lengths (N=50) and perithecia heights (N= 50) were measured using ImageJ (Schneider et al., 2012).

### *Fusarium graminearum* genetic manipulations

The *FgKnr4* gene was deleted through split marker-mediated transformation targeted fungal replacement with the hygromycin by homologous recombination (Yu et al., 2004). *F. graminearum* gene deletion construct assembly and fungal transformation was preformed following methods outlined in King et al., 2017. Primers were designed for the fusion of the 5’ and 3’ constructs using the NEBbuilder® Assembly Tool v.1 (https://nebuilderv1.neb.com/). Using the Gibson Master Mix (New England Biolabs, UK) the paired split marker fragments were ligated into the pGEM® -T Easy Vector (Promega, UK) then transformed into DH5α competent *Escherichia coli* (C2987H, New England Biolabs, UK) following standard manufacturer protocol. Diagnostic PCRs done using DreamTaq polymerase (ThermoFisher, UK) and standard cycling conditions. For the single gene deletion, three separate transformants two diagnostic PCRs detect the presence of the replacement cassette flanks (P3-4,P5-6) and the absence of the wild-type gene (P1-2) **(Figure 6 – figure supplement 1 A-B**). Complementation was performed following the protocol developed by Darino et al. (2024). Diagnostic PCRs for the complemented strains involved amplification of insertion cassette flanks (P7-8; P9-10), absence of short 868 bp empty intragenic locus amplicon (P11-P12), and test for heterozygosity of geneticin gene (P13-P14) **(Figure 6 – figure supplement 2 A-B**. Full primer list available in Supplementary File 3.

### Wheat host inoculation

The susceptible spring wheat (*T. aestivum*) cultivar, Bobwhite, was grown to anthesis. The 5th and 6th spikelets from the top of the wheat spike were inoculated on both sides using 5 μl of 5 x 10^5^ conidia/ml. Each treatment included 10 separate wheat plants (N=10). After inoculation, plants were kept in a high humidity chamber for 48 h in the dark. Disease progression was documented every two days by scoring the number of bleached spikelets. At 15 dpi wheat spikelet tissue and adjacent rachis internode was separated, frozen in liquid nitrogen, and ground to form a fine powder. The presence of DON mycotoxin was assessed using the Deoxynivalenol (DON) Plate Kit (Cat. 20-0016, Beacon Analytical Systems Inc., USA) following standard protocol. Experiment was replicated with three biological replicates per treatment (N=3). All plate images were taken using an Olympus OM-D camera using a 60mm ED M.Zuiko macro lens.

For resin dissection microscopy wheat cv. Bobwhite was inoculated 7^th^ and 8^th^ true spikelets from base inoculated each side w/ 5×10^5^ spores /ml in dH2O. After inoculation, plants were kept in a high humidity chamber for 48 h in the dark. Lemma tissues were excised from infected spikelets at 7 dpi, fixed in a 4% paraformaldehyde, 2.5% glutaraldehyde solution with 0.05M Sorensen’s phosphate buffer (NaH2PO4:Na2HPO4, pH 7.0). Samples then underwent 3 further buffer washes, a subsequent ethanol dehydration protocol (0-100% EtOH) over 48hrs and LR White resin (TAAB) infiltration diluted with dry ethanol at increasing ratios (1:4, 2:3, 3:2, 4:1, 100%). Samples were inserted into capsules (TAAB) and resin polymerised at 60°C for 16 hours in a nitrogen oven (TAAB). Ultra-thin 1µm sections of samples were cut on an ultramicrotome (Reichert-Jung, Ultracut) with glass knives, placed onto glass polysine slides (Sigma Aldrich, UK), dried at 70°C, stained with 0.1% (w/v) Toluidine Blue O and mounted in DPX mounting medium (Fisher Scientific). Stained sections were imaged on a Zeiss Axioimager (Axiocam 512 color, Zeiss, Jena, Germany) light microscope with brightfield illumination.

### Gene expression of module W12 genes

Bobwhite wheat plants were point inoculated at anthesis with either wild-type PH-1, *ΔFgtri5* or water only (Mock) following the protocol outlined in Dilks et al., (2019). Each experimental condition was replicated in triplicate, with each replicate deriving from three pooled independent wheat spikes. Tissues from rachis internodes 1 and 2 were sampled and frozen in liquid nitrogen at 3 dpi. Frozen samples were ground and RNA was extracted using the Monarch® Total RNA Miniprep Kit (NEB, UK). Equal amounts of RNA was used to synthesise cDNA with Revertaid cDNA synthesis kit (ThermoScientific, UK). PowerTrack™ SYBR Green Master Mix (ThermoScientific, UK) was used for qPCR. Each biological replicate included three technical replicates.

### Western blot

A 200 ul aliquot of a *F. graminearum* spore solution (1 x 10^6^ spores/ml) was added to 10 ml potato dextrose broth (PDB) at 27 °C. Calcofluor white was added to a concentration of 200 µg/ml after 24 h of incubation at 180 rpm. Twenty-four hours after the addition of the stress, mycelium was harvested, flash frozen and freeze dried To lyse the samples Y-PER Yeast Protein Extraction Reagent (ThermoScientific, UK) was added to the freeze-dried samples at 1.5 ml per 150 mg tissue, alongside Protease Inhibitor Cocktail (100x) (ThermoScientific, UK). Samples were lysed using the FastPrep-24™machine for 20s (MP Biomedical, USA). The supernatant was mixed with 5xSDS loading buffer (National Diagnostics, USA).

Equal amounts of protein (60 µg) were resolved on 8% SDS-PAGE gels (Mini-PROTEAN, Bio-Rad, UK) and transferred on to a nitrocellulose membrane. Immunoblots were performed by standard procedures using the Phospho-p44/42 MAPK (Erk1/2) (cat. #4370) and p44/42 MAPK (Erk1/2) (cat. #9102S) (Cell Signalling Technologies, USA) antibodies at their specified dilutions. The blots were developed using ECL Plus Western Blotting Detection Kit and images were acquired using Odyssey Imaging System (LI-COR Biosciences Ltd, Cambridge, UK).

### Microscopic examination of cell wall

Spores were induced by plating 200 µl of frozen spores (1 x 10^6^) PDA and incubating plates in for 3 days. For conventional transmission electron microscopy (TEM), fresh spores were harvested the same day from the PDA plates and pellets were fixed in a mixture of 2.5% glutaraldehyde and 4% Paraformaldehyde in Sorenson’s buffer (SB) at pH 7.2 overnight at 4°C. The samples were rinsed in SB and post fixed in 1% osmium tetroxide for 60 min at room temperature. The samples were dehydrated for 10 min per step into increasing concentrations of alcohol (30%, 50%, 70%, 90% and final 100%×3). Subsequently, the pure alcohol was replaced with propylene oxide, and the specimens were infiltrated with increasing concentrations (25%, 50%, 75%, and 100%) of Spurr resin mixed with propylene oxide for a minimum of 2 hr per step. The samples were embedded in pure, fresh Spurr resin and polymerised at 60 °C for 24 hr. Ultrathin sections (70 nm) were cut using an ultramicrotome (Leica UC7, Germany) and post-stained, first with uranyless for 1 min and then with Reynolds lead citrate for 2 min at room temperature, prior to observation using a Transmission Electron Microscope (Jeol 2100plus, UK) operated at 200 kV. *F. graminearum* spore solution (1 x 10^6^ spores/ml) was stained with Wheat Germ Agglutinin, Alexa Fluor™ 488 Conjugate (WGA) (10 μg/ml) for 10 minutes each. Samples were washed three times in sterile distilled water after staining. A ZEISS 780 Confocal Laser Scanning Microscope (ZEISS, Germany) was used to image spores.

### Phylogenetic tree construction

Eggnogmapper-v5 (Huerta-Cepas et al., 2019) was used to map *FgKnr4* to the eggnog Orthologue Group (OG) ENOG502QTAZ and generate the phylogenetic tree. The tree was visualised and annotated using the interactive Tree of Life (iTOL) software (Letunic and Bork, 2024).

### Functional characterisation of the *Knr4* orthologue in *Z. tritici*

Separate analyses using Orthologous Matrix (OMA) (Altenhoff et al., 2021) and Eggnogmapper (Huerta-Cepas et al., 2019) identified a single orthologous sequence in the genome of *Z. tritici* isolate IPO323 (https://fungi.ensembl.org/Zymoseptoria_tritici/Info/Index) (Goodwin et al., 2011). The gene has a Rothamsted gene model Id of ZtritIPO323_04g12347 (King et al., 2017; Chen et al., 2023) and is present on Chromosome 8 at start position 230142 bp. This maps to Mycgr3P105330 in the current genome call on Joint Genome Institute (JGI) Mycocosm (Goodwin et al., 2011).

Agrobacterium-mediated fungal transformation (Motteram et al., 2011) was performed to generate a series of independent gene disruption mutants of *ZtKnr4*. Flanking sequences and the hygromycin resistance gene were amplified from either genomic DNA or from plasmid pCHYG and using Phusion polymerase (NEB, UK). Fragments were gel purified using QIAquick Gel Extraction Kit (QIAGEN, UK) and assembled into the backbone (Kpn1 and BamH1 digested) of pCHYG by Gibson Assembly (NEB, UK). The resulting plasmids were transformed into Agrobacterium strain AgL1 and fungal transformation of isolate IPO323 was performed as per standard protocols (Motteram et al., 2011). Positive transformants containing a disrupted *ZtKnr4* gene were identified by diagnostic PCR **(Figure 8 – figure supplement 1**). Complementation of validated *ZtKnr4* mutants was performed through Agrobacterium-mediated transformation with plasmid pCGEN (digested EcoR1 and Kpn1) containing the native gene plus 1 kb upstream (5’) and 300 bp (3’) downstream genomic DNA, amplified by Phusion PCR (NEB, UK).

Attached leaf virulence assays were performed as per standard protocols (Keon et al., 2007) on wheat cultivar Riband. Leaf blades (N = 3) were inoculated with spore suspensions of 1 x 10^6^ spores / ml in sterile water + 0.05% v:v Tween 20. Final disease assessments were made 21 days after inoculation. *In vitro* hyphal growth assays were performed following droplet inoculation of spore suspensions onto 1% Tap Water Agar (TWA) plates. Hyphal growth morphologies were determined by light microscopy and / or photography 10 days after inoculation. Calcofluor white sensitivity assays were performed to ascertain changes in cell wall strength. For this, spore suspensions were inoculated onto YPD agar (Formedium, UK) plates (control) and onto YPD agar plates containing 200 μg / ml calcofluor white. Plates were incubated at RT for 8 days and then growth was monitored and recorded by photography. Images of *ZtKnr4 in planta* and *in vitro* experiments were taken with a Nikon D3200 camera.

## Data availability

Full lists of all genes clustered into modules is available on https://github.com/erikakroll/Fusarium-wheat_WGCNA. This includes comma separated value (CSV) files for all genes in each module for both fungal and wheat modules, which are annotated with Module Membership (MM) values, mean FPKM values, InterPro annotation, Gene Ontology annotation, and Trait Ontology annotation. Text documents containing module eigengene values and gene module assignments are also available on the repository.

## Author’s contributions

E.K conducted the experiments and wrote the manuscript. N.A.B, M.U, and K.H.K provided project oversight, experimental design and manuscript planning, development, and revisions. R.A helped with experimental design and data analysis.

C.B and J.R generated ZtKnr4 mutant and completed associated characterisation experiments. A.M.U embedded, sectioned, and imaged samples for TEM analysis.

V.A undertook the resin embedding, sectioning, and imaging.

---

**Figure 2 – figure supplement 1.**
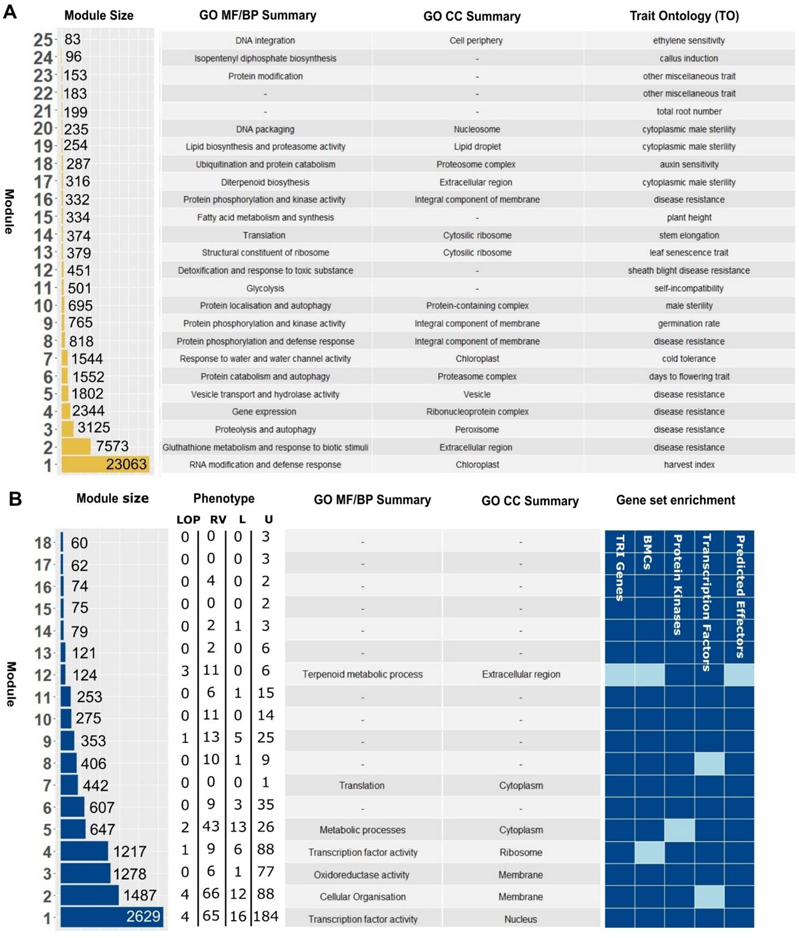
Network Summary. **A.** Summary of all modules in the wheat network, including module size (number of genes), Gene Ontology (GO) and Trait Ontology (TO) enrichment summaries. B. Summary of all modules in the fungal network, including modules size, Gene Ontology (GO) enrichment summaries and Gene Set Enrichment Analysis (GSEA). The number of genes with different phenotypes in PHI-base are depicted, with LOP, RV, L and U denoting different PHI-base phenotypes (LOP = Loss of pathogenicity; RV = Reduced virulence; L = Lethal; U = Unaffected pathogenicity) **(Urban et al., 2022**).

**Figure 2 – figure supplement 2.**
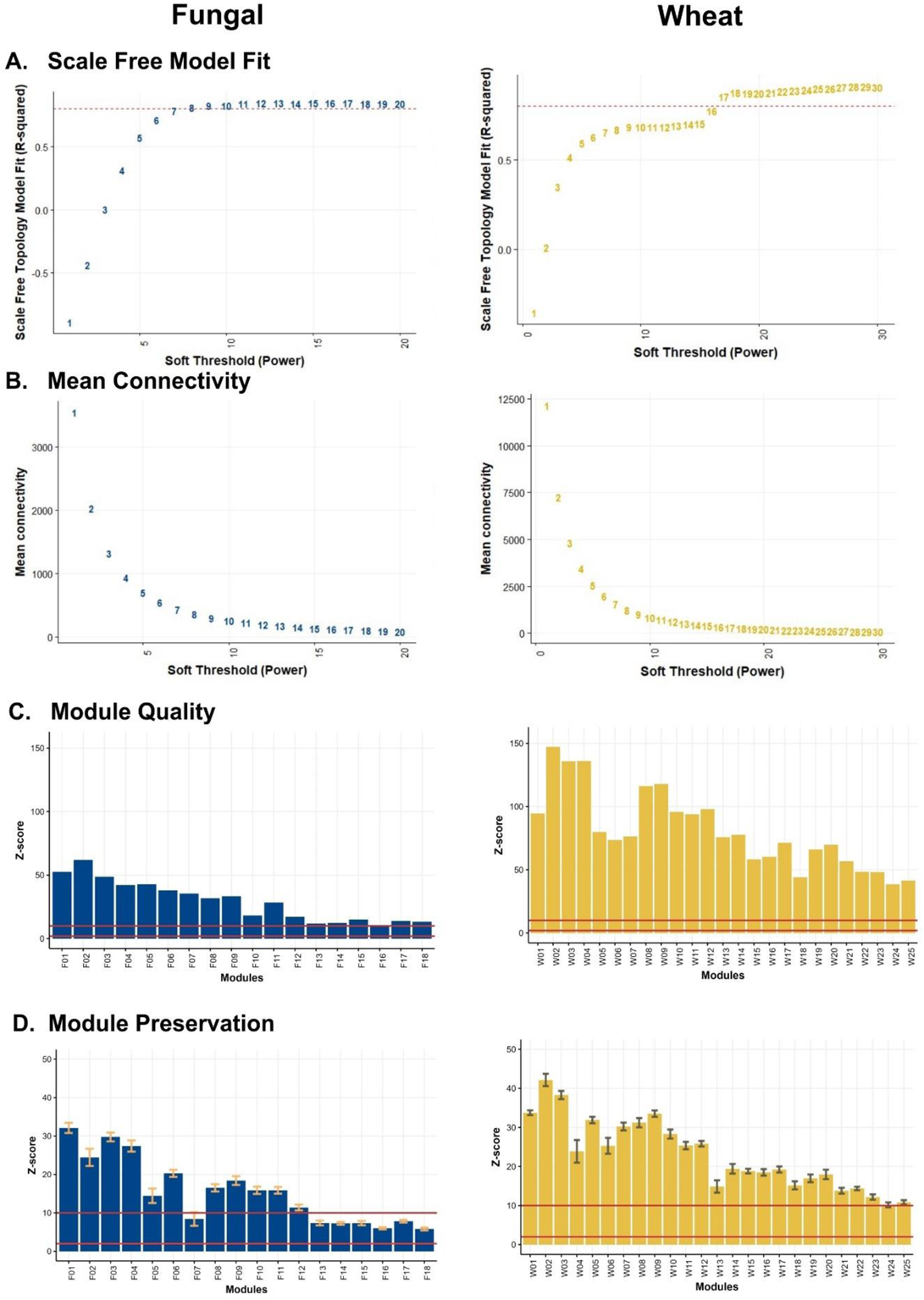
**Network statistics. A**. Strength of correlation of network model (R-squared value) to scale free model at different soft thresholding powers. Dotted red line is at an R-squared value of 0.80, the threshold needed for generating a WGCNA network. **B.** Mean connectivity of genes in each network at different soft thresholding powers. A low mean connectivity is desired to meet the scale free network criteria. **C.** Module quality across all modules as determined by a Z-score calculation. Solid red lines at minimum quality (Z = 2) and high quality scores (Z = 10). **D.** Module preservation as determined by Z-score calculation against 50 random test networks. Solid red lines at minimum preservation (Z = 2) and high preservation scores (Z = 10).

**Figure 4 – figure supplement 1.**
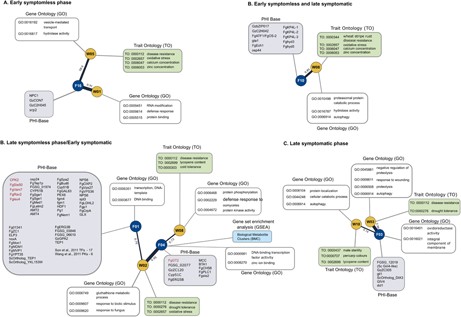
Annotation of stage-specific modules. Fungal modules (F) and wheat modules (W) depicted with significant enrichment annotations. Fungal modules are additionally annotated with known phenotypes from PHI-base. Genes listed in red in the PHI-base annotation (grey box) exhibit a loss of pathogenicity phenotype when deleted, while the remaining genes display a reduced virulence phenotype when deleted. Plots are separated by modules with highest expression in a given stage of infection, namely **A. Early symptomless, B. Early symptomless and late symptomatic, C. Late symptomless/Early symptomatic, and D. Late symptomatic.**

**Figure 5 – supplement 1.**
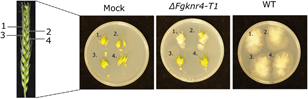
Surface sterilisation of dissected wheat floral tissue. . Dissection at 15 dpi of wheat spikes followed by separation of infected wheat spikelet and rachis tissues and subsequent plating onto synthetic nutrient agar (SNA) separated at 15 dpi. Plate images taken 3 days later.

**Figure 6 – figure supplement 1.**
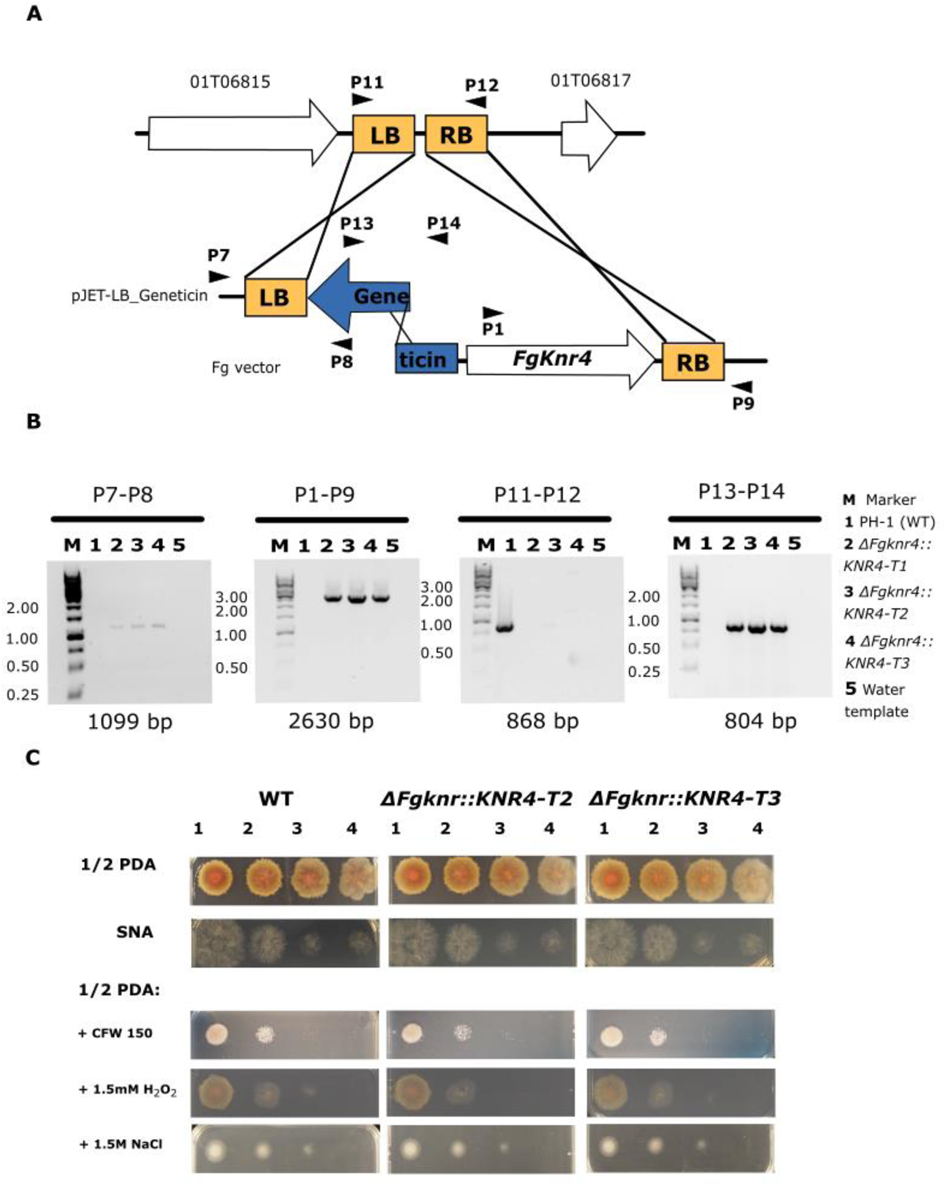
*FgKnr4* single gene deletion strategy and characterisation of additional transformants. **A.** Schematic for the hygromycin split marker deletion strategy including diagnostic primer locations (P1-6). **B.** Diagnostic PCR with primer sets depicted in panel A. PCR samples were separated on 7.5 % agarose gel with a 1 kb DNA ladder. The expected amplicon size is written below the corresponding gel image. **C.** Dilution series of wild-type (WT) and additional *ΔFgknr4* transformants (*T2* and *T3*) on Synthetic Nutrient Agar (SNA) and half-strength Potato Dextrose Agar (0.5 PDA) with and without the addition of single stresses. The dilution series begins at 1: 1 x 10^6^ and continues with 10-fold dilutions (2: 1/10, 3: 1/100, and 4: 1/100). Images taken after 3 days.

**Figure 6 – figure supplement 2.**
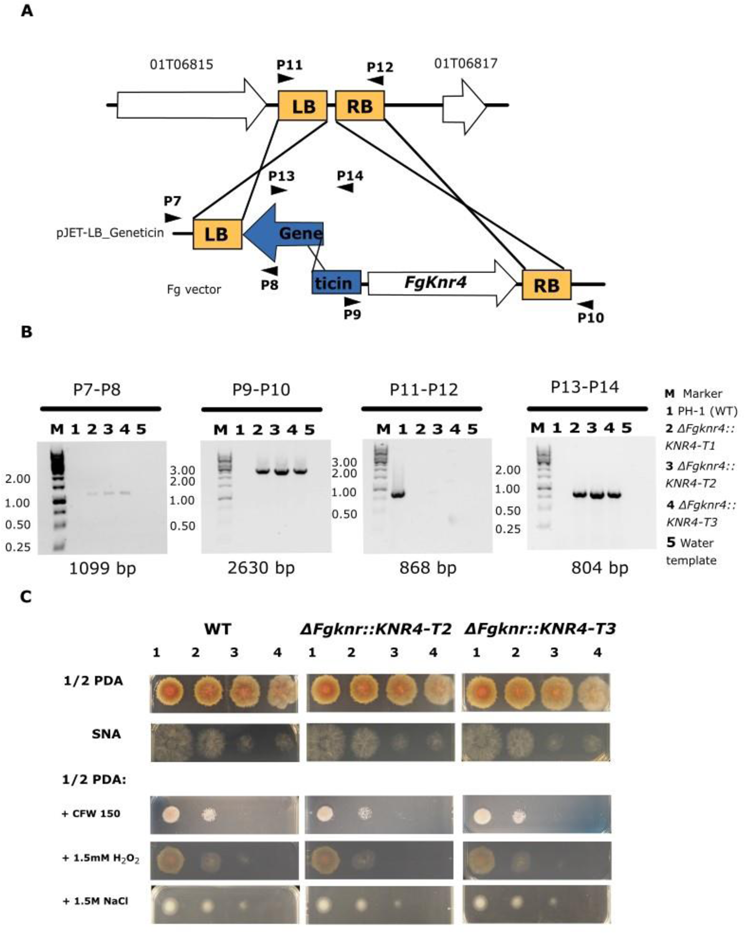
*ΔFgknr4-T1* complementation strategy and characterisation of additional transformants. **A.** Schematic of gene complementation into the *Fg* transformation locus (Darino et al. (2024), including diagnostic primer locations (P7-P14). **B.** Diagnostic PCR with primer sets depicted in panel A. PCR samples were separated on 7.5 % agarose gel with a 1 kb DNA ladder. Expected amplicon size is written below the corresponding gel image. **C.** Dilution series of wild-type (WT) and additional *ΔFgknr4::KNR4* transformants (*T2* and *T3*) on Synthetic Nutrient Agar (SNA) and half-strength Potato Dextrose Agar (0.5 PDA) with and without the addition of single stresses. The dilution series begins at 1: 1 x 10^6^ and continues with 10-fold dilutions (2: 1/10, 3: 1/100, and 4: 1/100). Image taken after 3 days.

**Figure 6 – figure supplement 3.**
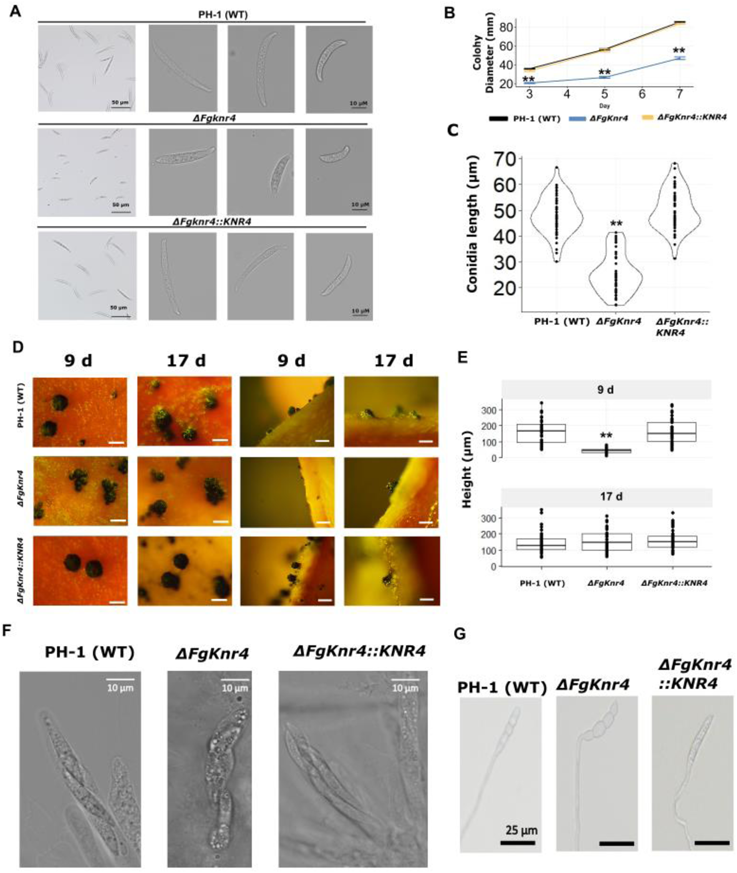
Characterisation of growth rate, conidial size and ascospore production in *ΔFgknr4* and complemented strains. A. Decreased condial size observed in *ΔFgknr4.* Single conidial images to represent long, middle length, and short conidia across strains. B. Mean colony diameter of wild-type (WT), *ΔFgknr4*, and *ΔFgknr4::KNR4* grown on Potato Dextrose Agar (PDA) (N=5). C. Distribution of conidial length from N = 50 spores for wild-type (WT), *ΔFgknr4*, and *ΔFgknr4::KNR4* strains. D. Representative perithecia images taken after perithecia induction in carrot agar medium using wild-type (WT), *ΔFgknr4*, and *ΔFgknr4::KNR4* strains. Images taken from above (left panels) and from agar sections placed on slides (right panels) on day 9 and day 17. Scale bar = 500 µm. F. Ascospores in intact ascus produced by wild-type (WT), *ΔFgknr4* or *ΔFgknr4::KNR4* strains. Scale bar = 10 µM. G. Ascospores obtained from squashed perithecia of wild-type (WT), *ΔFgknr4* or *ΔFgknr4::KNR4* strains are viable and form germ tubes. Scale bar = 25 µm. Significance is denoted as **= *p* ≤ 0.01. Significance was determined by a one-way ANOVA followed by Tukey HSD correction.

**Supplementary Figure 6 – figure supplement 4.**
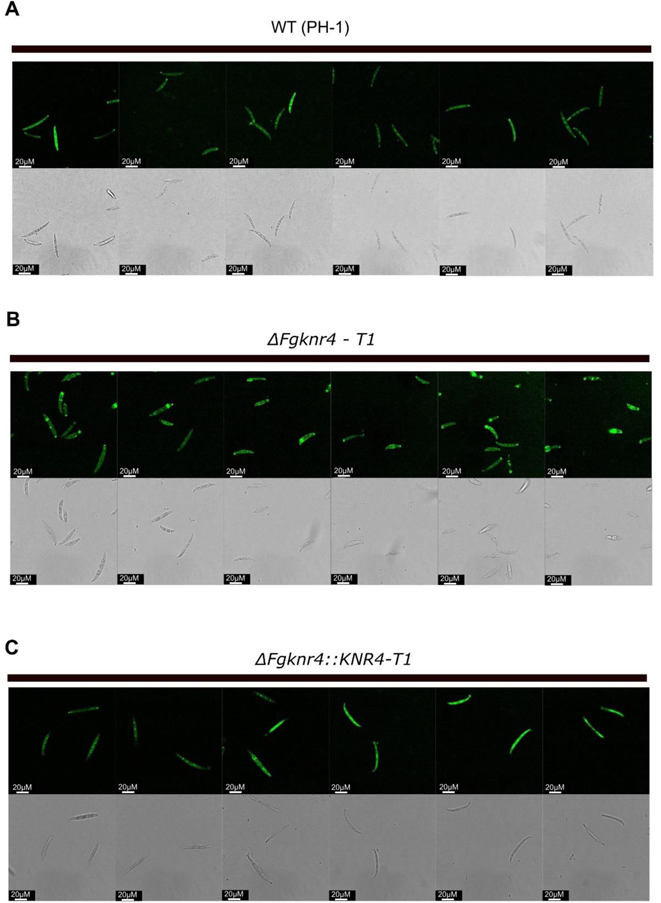
Additional fluorescent microscopy images of irregular chitin distribution in *ΔFgknr4* conidia. . Visualisation of chitin-stained conidia with Wheat Germ Agglutinin Alexa Fluor™ 488 Conjugate (WGA) in wild-type (WT) (A), *ΔFgknr4* (B) and *ΔFgknr4::KNR4* (C).

**Figure 6 – figure supplement 5.**
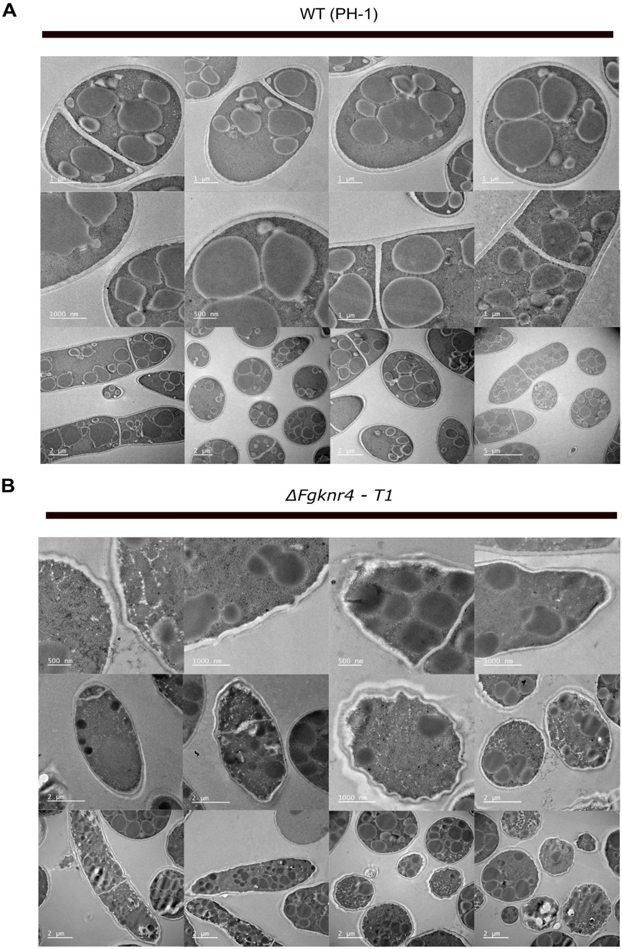
Additional TEM images of abnormal cell wall morphology in *ΔFgknr4* conidia. TEM imaging of wild-type (**A**) and *ΔFgknr4* (**B**) conidia, showing differences in cell wall structure and different magnifications.

**Figure 8 – figure supplement 1.**
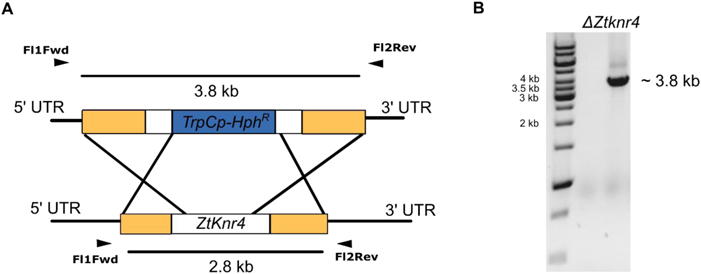
*ΔZtknr4* diagnostic PCR and disruption deletion strategy. **A.** Hygromycin (*Hph^R^*) replacement cassette inserts at *ZtKnr4* locus through homologues recombination via homologous flanks (yellow and white). **B.** Diagnostic PCR demonstrating presence of large insertion fragment in *ΔZtknr4* transformant using Fl1Fwd and Fl2Rev primers.

**Supplementary File 1. Network module sizes and gene module assignments.** Spreadsheet containing sizes of all modules in fungal and wheat networks. ‘Fungal module assignments’ and ‘Wheat module assignments’ tabs contain a column with all fungal IDs (RRES v.5 PH-1) or wheat IDs (Column A = IWGSC RefSeq v2.1; Column B = IWGSC RefSeq v1.1) with an adjacent column denoting which module they are clustered in.

**Supplementary File 2.** *F. graminearum* genes with known phenotypes with the PHI-base database (www.PHI-base.org) in each fungal module. Table provides RRES v5 gene ID, PHI identifier ID from PHI-base, Uniprot protein ID, gene function, mutant phenotype, author reference, and year published.

**Supplementary File 3. Primer list.** Primers used to generate mutant and complemented strains.

**Table S1.**
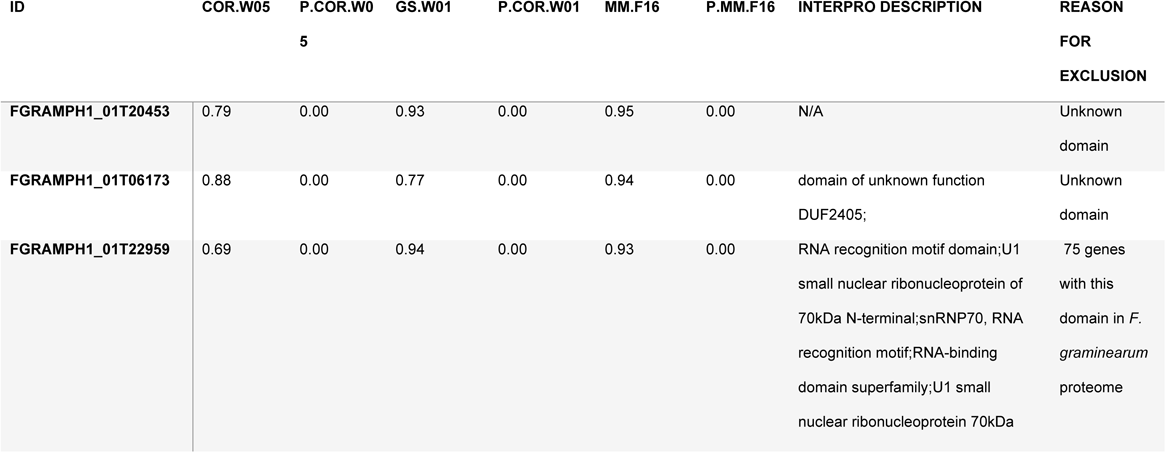
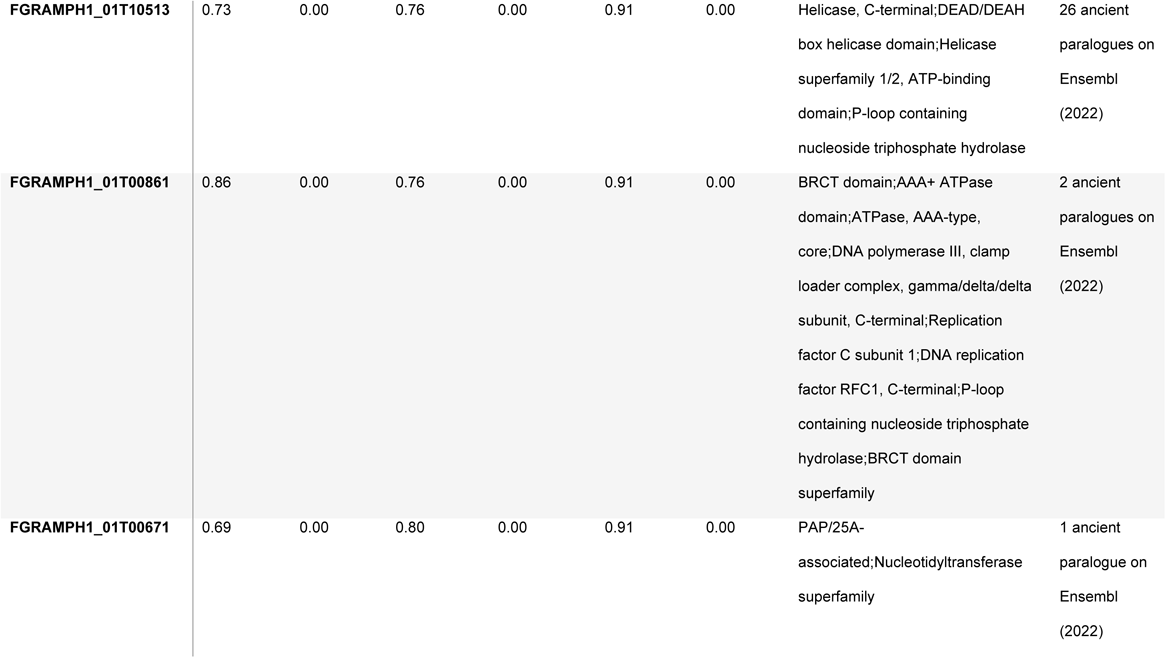
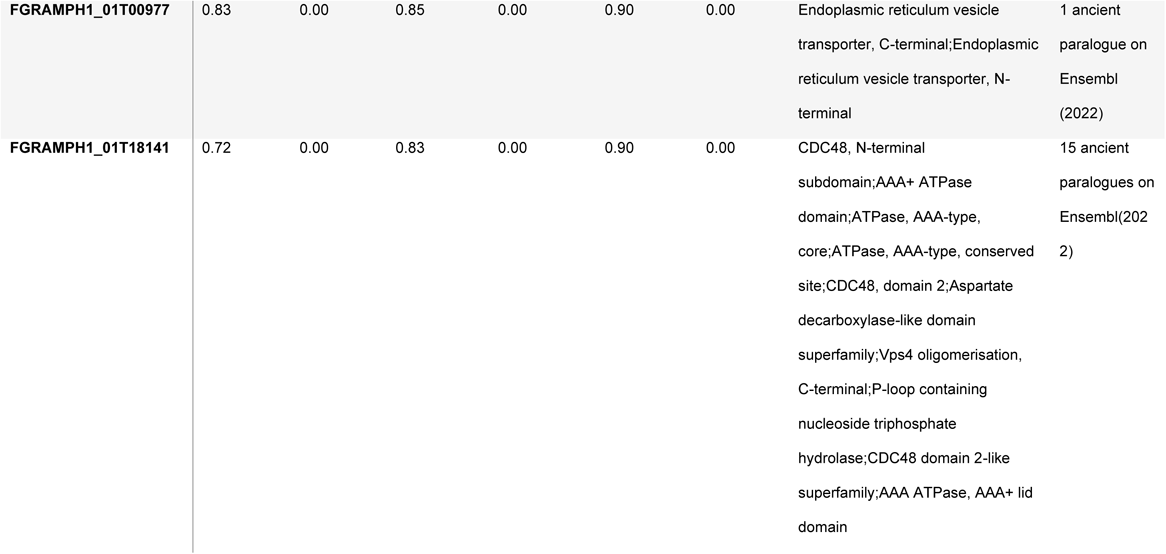
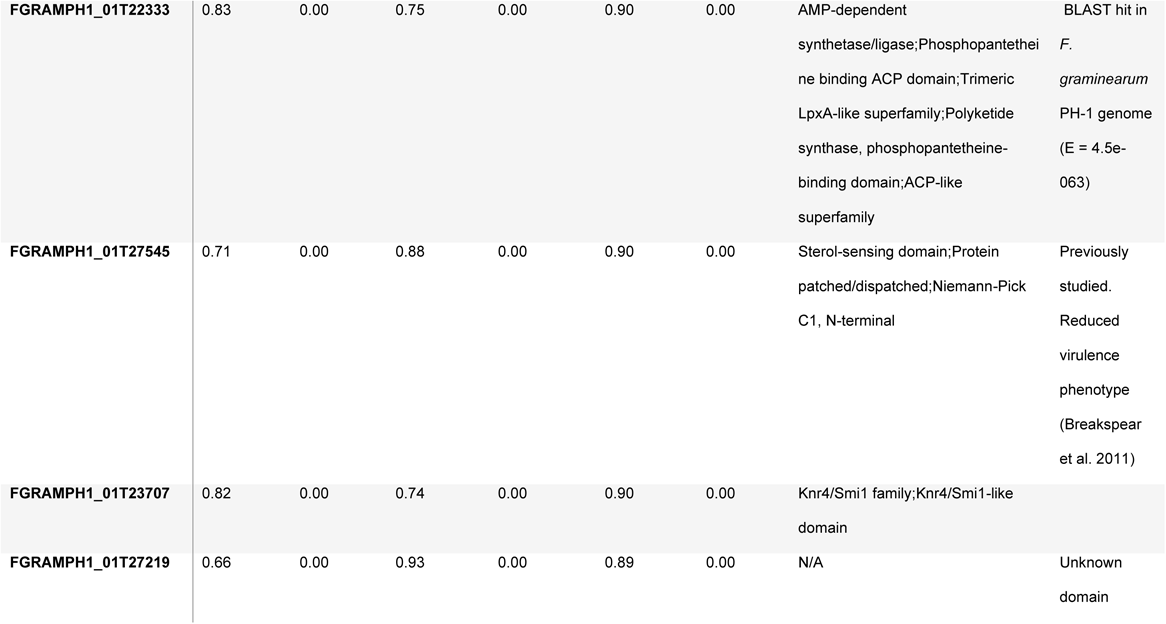
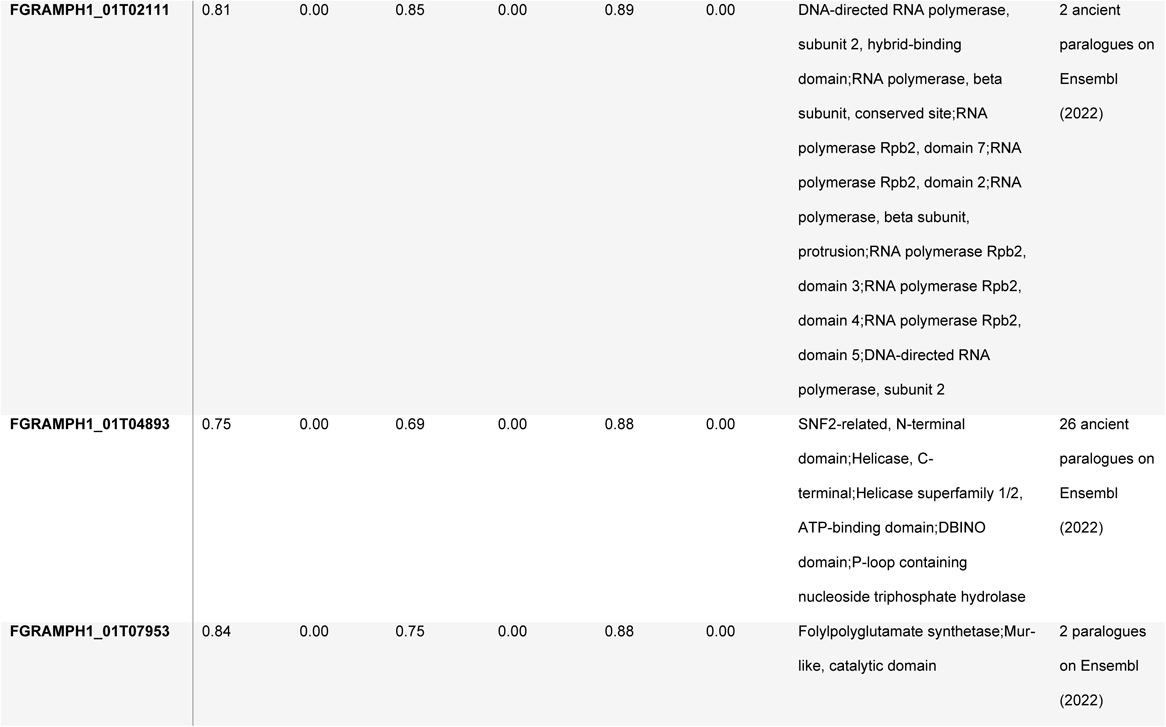
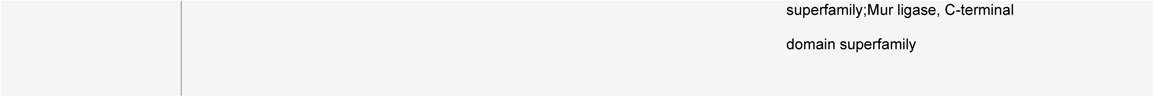
Candidate gene selection in fungal module F16. Table provides details on the 15 candidates within module F16 with the highest module membership (MM) and reason for exclusion from further functional characterisation analysis. This table includes the MM score and associated *p-*values (p.MM), as well as correlation strength to corresponding wheat modules (Cor) and *p-*values (p.Cor).

